# Forensic microbial system for high-resolution object provenance

**DOI:** 10.1101/2020.03.14.990804

**Authors:** Jason Qian, Zhi-xiang Lu, Christopher P. Mancuso, Han-Ying Jhuang, Rocío del Carmen Barajas-Ornelas, Sarah Boswell, Fernando H. Ramírez-Guadiana, Victoria Jones, Akhila Sonti, Kole Sedlack, Lior Artzi, Giyoung Jung, Mohammad Arammash, Michael Melfi, Lorena Lyon, Siân V. Owen, Michael Baym, Ahmad S. Khalil, Pamela Silver, David Rudner, Michael Springer

## Abstract

Mapping where an object has been is a fundamental challenge for human health, commerce, and food safety. Location-specific microbes offers the potential to cheaply and sensitively determine object provenance. We created a synthetic, scalable microbial spore system that identifies object provenance in under one hour at meter-scale resolution and near single spore sensitivity, and that can be safely introduced into and recovered from the environment. This system solves the key challenges in object provenance: persistence in the environment, scalability, rapid and facile decoding, and biocontainment. Our system is compatible with SHERLOCK, facilitating its implementation in a wide range of applications.

## Main text

Globalization of supply chains has dramatically complicated the process of determining the origin of agricultural products and manufactured goods. Determining the origin of these objects can be critical, as in cases of foodborne illness, but current labelling technologies can be prohibitively labor-intensive and are easy to remove or replace (*1*). Similarly, law enforcement could benefit from tools that label unknown persons or objects passing through a location of interest as a complement to fingerprinting and video surveillance (*2*). Microbial communities offer a potential alternative to standard approaches of labeling. Any object placed in and interacting with a particular environment gradually adopts the naturally occurring microbes present in that environment (*3,4*); thus, it has been suggested that the microbial composition of an object could be used to determine object provenance (*5*), although if whether transfer can occur in a relevant timescale is unclear. However, the composition of microbial communities in different areas are not reliably large or stable enough to uniquely identify specific locations; moreover, using natural microbes requires extensive, expensive, and time-consuming mapping of natural environments.

To circumvent these challenges, we propose the deliberate introduction and use of synthetic microbes harboring barcodes that uniquely identify locations of interest (e.g. food production areas). These microbes would offer a sensitive, inexpensive, and safe way to map object provenance provided that several important criteria are met, including: 1) the synthetic microbes must persist in the environment and reliably label objects that pass through it; 2) the microbes must be contained, preventing cross-contamination and adverse ecological effects; and 3) the encoding and decoding of information about the provenance of the object must be rapid, sensitive and specific. Similar barcoding approaches have been explored previously to model pathogen transmission (*6,7*), but did not explicitly address those challenges. Here we report the FMS (<u>F</u>orensic <u>M</u>icrobial <u>S</u>pores) system, a scalable, safe and sensitive system that uses DNA-barcoded microbial spore communities to permit the determination of object provenance (**Fig. 1A**).

**Fig. 1.**
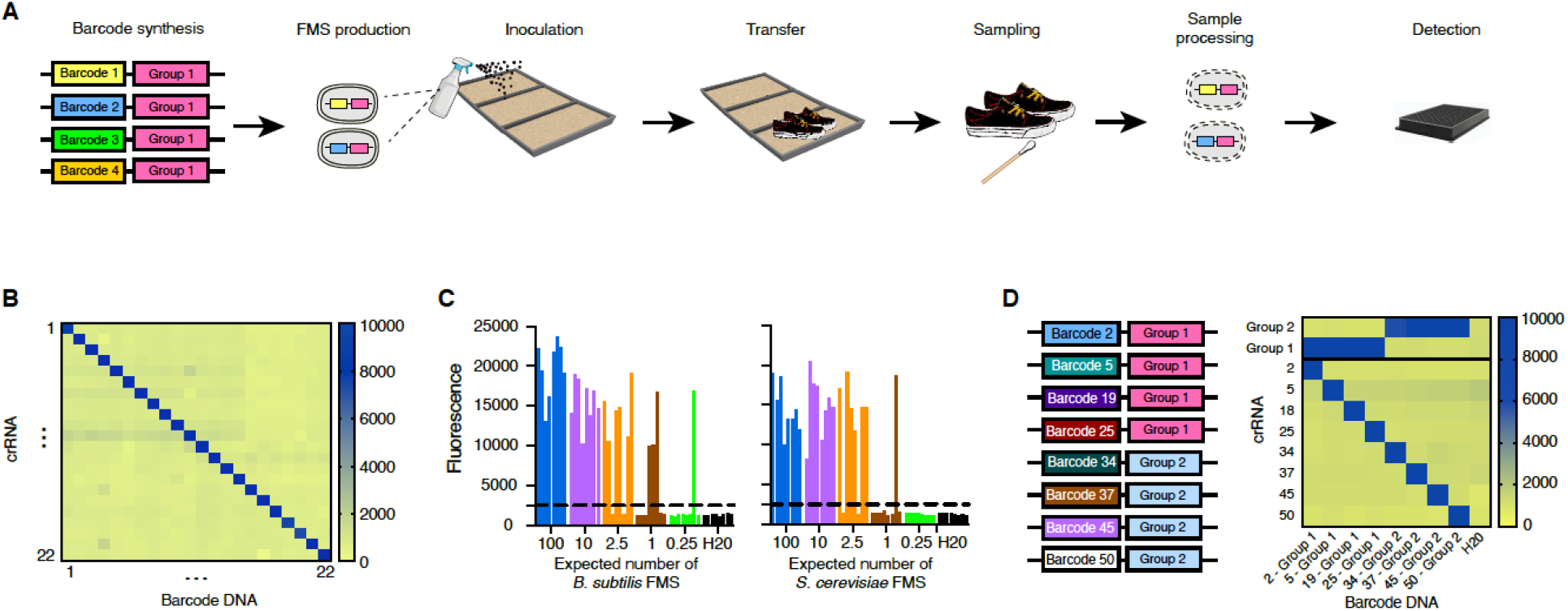
Forensic Microbial Spores can be specifically and sensitively detected. **(A)** Schematic of the Forensic Microbial Spores (FMS) application and detection pipeline. **(B)** Heatmap showing endpoint fluorescence values from *in vitro* SHERLOCK reactions of all combinations of 22 barcodes and 22 crRNAs assessing specificity of each barcode-crRNA pair. **(C)** Detection limit of various *B. subtilis* and *S. cerevisiae* FMS by SHERLOCK (each of the 8 biological replicates for each spore concentration are shown). Spore numbers are calculated on a per reaction basis. **(D)** Heatmap showing endpoint fluorescence values from *in vitro* SHERLOCK reactions testing specificity of 4 barcodes for group 1 crRNA and 4 barcodes for group 2 crRNA as detected by either unique or group crRNA.

The FMS system leverages the natural ability of spores to persist for long periods in the environment without growth (*8*). We designed unique DNA barcodes and integrated them into the genomes of *Bacillus subtilis* (*B. subtilis*) and *Saccharomyces cerevisiae* (*S. cerevisiae*) spores, creating a set of barcoded FMS that can be used combinatorially to provide a nearly infinite set of unique identification codes. The FMS can be: 1) manufactured at scale using standard cloning and culturing techniques; 2) applied to surfaces by spraying; and 3) depending on the surface colonized, efficiently transferred to objects that come into contact with the colonized surface. To identify barcodes, FMS sampled from objects are lysed and can be decoded with a range of methods including SHERLOCK, a recombinase polymerase amplification (RPA) method coupled with a Cas13a-based nucleic acid detection assay (*9*), qPCR, and sequencing (**Fig. 1A and fig. S1A**).

The FMS do not impact the native environment into which they are applied. First, we used auxotrophic strains that require amino acid supplementation for growth. Second, we made the cells germination deficient. For *B. subtilis* spores, we deleted the genes encoding the germinant nutrient receptors and the genes that encode the cell wall lytic enzymes required to degrade the specialized spore cell wall. Incubation of >10^12^ spores made from these mutant strains showed they were unable to form colonies or grow in rich medium, and remained stable and non-germinating at room temperature for >3 months (**figs. S2, A and C**). For *S. cerevisiae*, we boiled spores for 30 minutes to heat-kill vegetative cells prior to application. Incubation of >10^8^ boiled spores on rich medium yielded no colonies (**figs. S2, B and D-F**). Finally, all antibiotic resistance cassettes used to generate the FMS were removed by site-specific recombination to prevent horizontal gene transfer of resistance genes to other organisms in the environment. After integration, the inserted sequence does not encode for any gene and should not confer any fitness advantage if horizontally transferred.

Multiple barcoded FMS can be applied then decoded simultaneously. We designed a series of tandem DNA barcodes, each with a Hamming distance >5 allowing more than 10^9^ possible unique barcodes. To test the specificity of our barcode design in a field-deployable system, we constructed 22 barcodes and their matching crRNAs and assayed all permutations *in vitro* by SHERLOCK. All 22 crRNAs clearly distinguished the correct barcode target (**Fig. 1B**). In order to scale the system, we devised a facile method to quickly screen a large number of barcodes and crRNAs in parallel to eliminate those with cross-reactivity or background; we validated and performed pooled *n-1 barcodes* RPA reactions *in vitro* with corresponding crRNA and water RPA controls testing 94 crRNA-barcode pairs, eliminating 17 for high background and 7 for cross-reactivity (**figs. S1, B and C**). To test sensitivity and specificity *in vivo*, we integrated 57 barcodes into *B. subtilis* and 11 into *S. cerevisiae*. We developed an efficient spore lysis protocol using heat and sodium hydroxide (**fig. S3**), which allowed us to achieve near single-spore resolution for detection using SHERLOCK (**Fig. 1C**). *In vivo* and *in vitro* specificity screenings of crRNA-barcode pairs gave similar results (**fig. S1C**). Additionally, our barcodes are tandemly designed with a group and a unique sequence (**fig. S1A**) to aid in high-throughput detection settings where only a small subset of samples contains an FMS of interest. The group sequence is compatible with field-deployable detections and can be used to determine whether an FMS of interest is present before using a second assay to uniquely identify the FMS (**Fig. 1D**). This two-step process solves the throughput limitations of field-deployable detection and lowers the costs of sequencing.

The FMS system is robust and can function in real-world environments. First, in ~1 m^2^-scale experiments (**fig. S4A and Table S7**), we used qPCR to detect and quantify FMS directly from surface samples or surface swabs. We found that FMS persisted on sand, soil, carpet, and wood surfaces for at least 3 months with little to no loss over time (**Fig. 2A and figs. S4B-C**). Notably, multiple tested perturbations (e.g. simulated wind, rain, vacuuming, or sweeping in **fig. S4A**) did not significantly reduce our ability to detect FMS from the surface. Second, we constructed a ~100 m^2^ indoor sandpit (**Fig. 2B and fig. S5**), colonized one region with FMS (**Fig. 2C**), and were able to readily detect the FMS for 3 months using SHERLOCK (**Fig. 2D**). Importantly, perturbations did not cause appreciable spreading to non-colonized areas (**Fig. 2D and fig. S6**), even a catastrophic disturbance where a fan fell into the pit displacing a significant amount of sand only spread the FMS several meters (**fig. S5**). This is consistent with low levels of re-aerosolization reported for other spore-forming *Bacilli* (*10*).

**Fig. 2.**
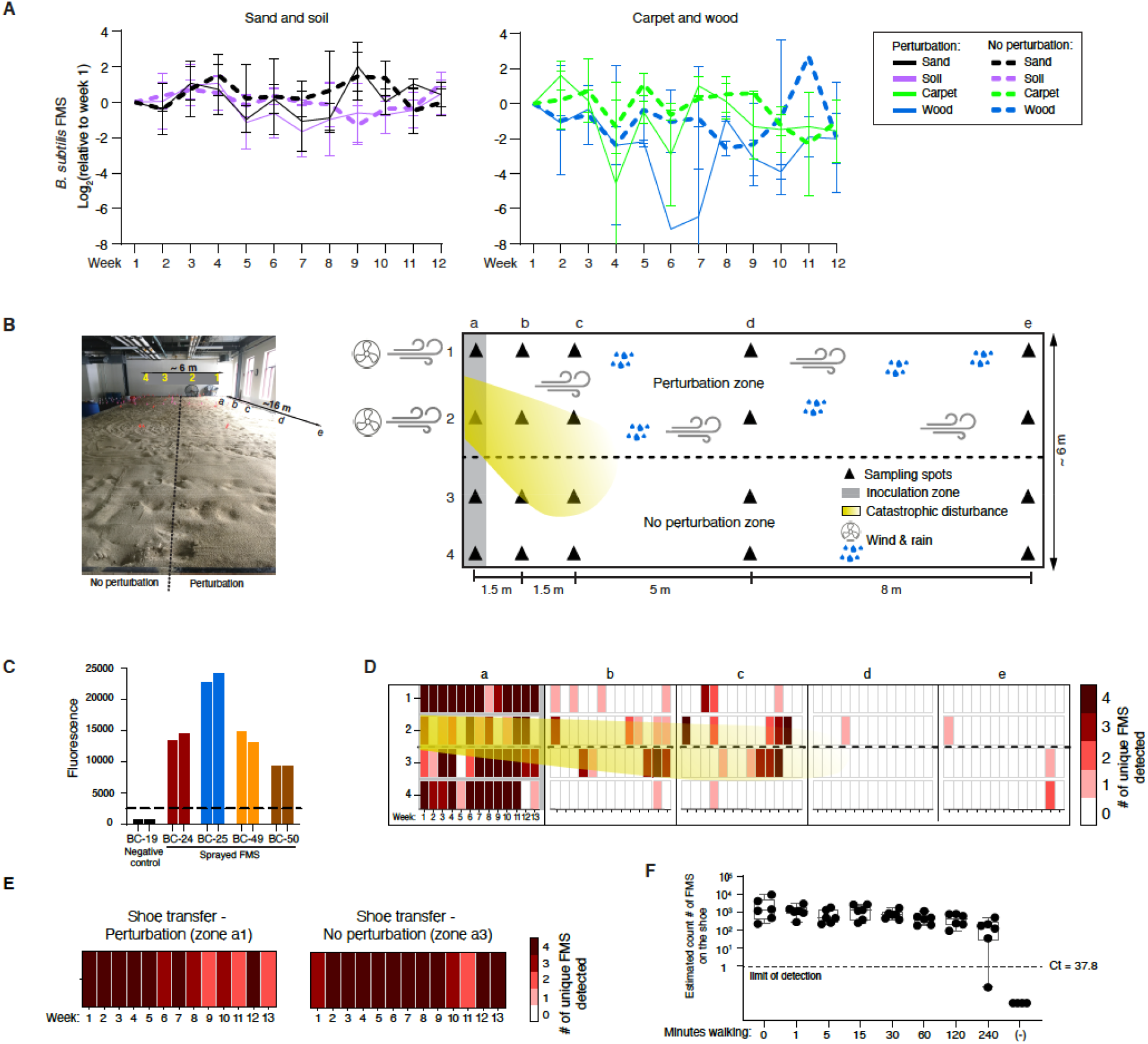
Persistence, transferability, and maintenance of FMS. (**A**) Barcoded FMS persisted on sand, soil, carpet, and wood over 3 months on ~1 m^2^ test surfaces performed in incubators; y-axis: *B. subtilis* FMS number relative to week 1 levels. (Bars represent standard deviations). Perturbation: simulated wind, rain, vacuuming, or sweeping. (**B)** Photograph of large-scale (~100 m^2^) and schematic of sandpit. *B. subtilis* BC-24 and BC-25 FMS, and *S. cerevisiae* BC-49 and BC-50 FMS were colonized in zone a, shaded in grey. The yellow shadow represents the area over which the top 2 inches of sand from a 1.5 m^2^ area of zone a was redistributed after a large fan fell over. **(C)** SHERLOCK signal from all 4 barcoded FMS used for large-scale colonization. Dashed line is the threshold for positive calls. (**D**) FMS persist in zone a and do not spread to zone d or e. Heatmap depicts the number of barcoded FMS (out of 4) detected by SHERLOCK. (**E**) FMS were transferred onto shoes by walking in the sandpit zone a, and were detected by SHERLOCK. **(F)** Abundance of BC-25 FMS on shoes after up to 240 minutes of walking on uncolonized areas; y-axis: FMS count based on qPCR standard curve.

The FMS can be transferred onto objects that pass through test environments. At ~1 m^2^ scale, FMS could be transferred onto rubber or wooden objects simply by placing them on the FMS-colonized surface for several seconds, yielding up to ~100 spores per microliter of reaction input (**fig. S4D**). At larger scale, FMS were reliably transferred onto shoes worn in the colonized sandpit (**Fig. 2E and fig. S7**). Furthermore, the FMS transferred onto shoes could be still detected on the shoes even after walking on uncolonized surfaces for several hours, though FMS counts decreased by 2-fold with 2 hours of walking as quantified by qPCR (**Fig. 2F and fig. S8**). We conclude that the FMS can persist in the environment without significant spreading; are transferable onto objects that pass through the environment; are retained on these objects; and can be sensitively and specifically detected using SHERLOCK.

The FMS system can be used to determine whether an object has passed through a specific environment. We colonized 12 quadrants of sand, each with 4 unique FMS (**Fig. 3A and fig. S9B**) and traversed them with a series of test objects. To mimic in-field deployment, we used a portable light source, an acrylic filter, and a mobile phone camera to image the SHERLOCK readout (**Fig. 3B and fig. S10B**). We successfully determined object provenance in >20 test objects with only a single false positive quadrant (**Fig. 3C and fig. S9**). Notably, provenance could be determined in the field in ~1 hour from sample collection. To evaluate the sensitivity and specificity of our system for determining an object’s trajectory, we tested more objects, changing the number of unique FMS colonized in each quadrant, as well as the surface material (i.e. soil, sand, carpet, and wood) (**figs. S9-11**). As expected, increasing the number of unique FMS used per quadrant improves the confidence levels of positive calls by adding redundancy to the decoding. While 1 or 2 unique FMS per quadrant did permit us to determine object trajectories, we found optimal results when using 4 unique FMS, i.e. a 0.6% (1/154) false positive rate and 0% (0/62) false negative rate. (**Fig. 3C and figs. S9-11**). Importantly, the FMS were positively detected on all objects regardless of which of the 4 colonized surfaces the objects passed through (**fig. S11**). More broadly, this experiment demonstrates that the FMS can be used to determine object provenance at meter-scale resolution, which would be extremely difficult to achieve using natural microbiome signatures (*11*).

**Fig. 3.**
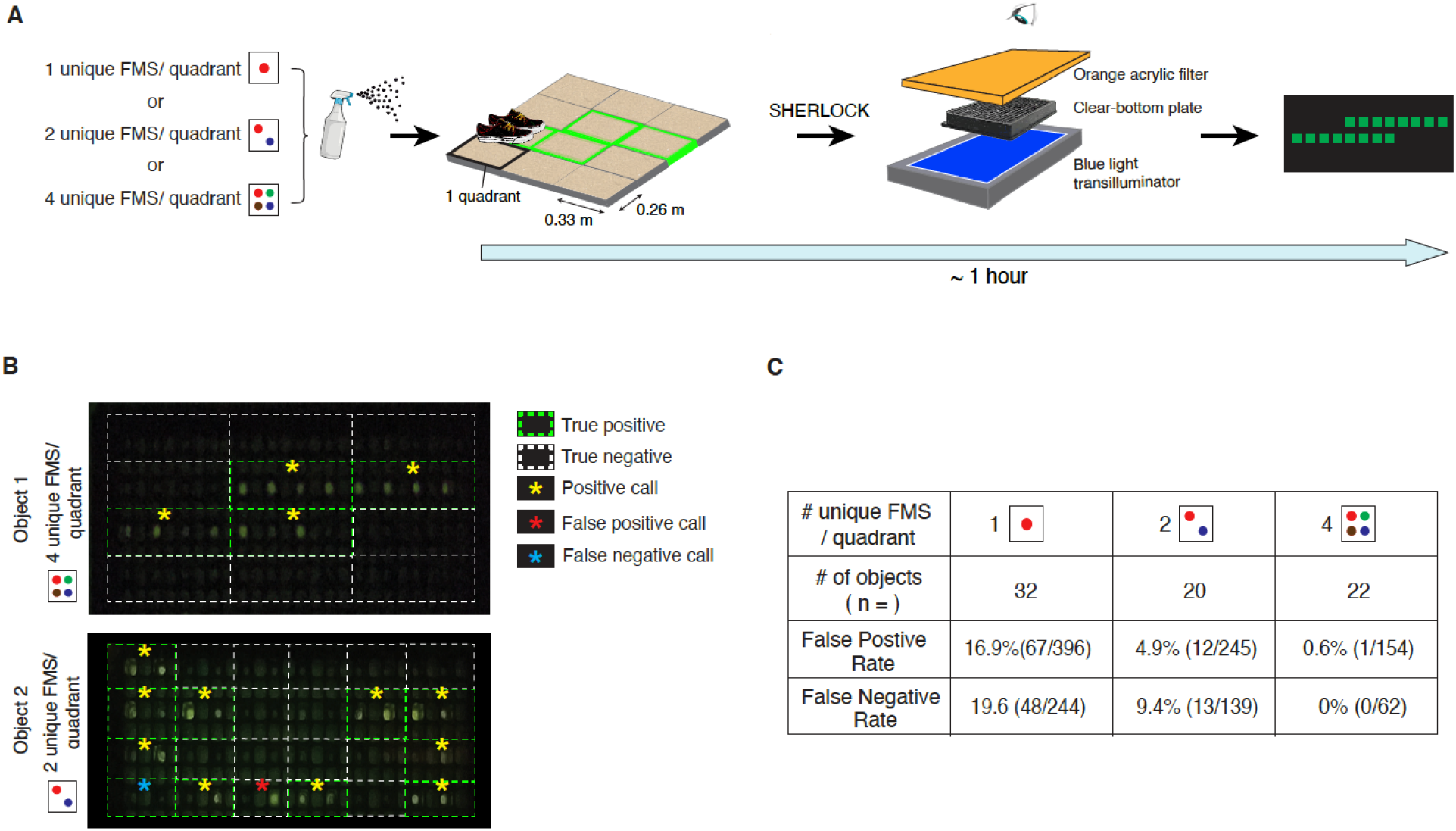
Determination of object provenance using FMS and field-deployable detection. **(A)** Schematic of experiment design and field-deployable detection method. Each quadrant was inoculated with 1, 2, or 4 unique barcoded FMS. Green outlines indicate the object trajectory through a subset of the quadrants. SHERLOCK reactions were imaged using a mobile phone camera to photograph the reaction plate through a filter, under portable blue light illumination. **(B)** Reaction plate images in which each region is overlaid with a yellow * to denote a positive call, a red * for false positive call, a blue * for false negative call, a green dashed outline for true positives, and a white dashed outline for true negatives. **(C)** Statistics for SHERLOCK trajectory predictions of objects traversing quadrants colonized with 1, 2, or 4 unique barcoded FMS/ quadrant.

The FMS system can determine food provenance. Foodborne illness is a global health issue with an estimated 48 million cases each year in the US alone (*12*). There is an urgent need for rapid methods for identifying the source of food contamination, as it often takes weeks and is costly with current approaches due to the complex modern market chain (*13*). *Bacillus thuringiensis* (*Bt*) is an FDA-approved biocide that is widely used in agriculture (*14*). To determine if *Bt* spores applied during farming could be detected in produce purchased from a market, we used qPCR to quantify the *Bt* cry1A gene. *Bt* was detected on all plants that were treated with *Bt* and not detected on plants that were not treated with *Bt* (**fig. S12**). In addition, we detected *Bt* spores in 19 out of 32 pieces of produce bought at various stores (**Fig. 4A and fig. S13, and Table S1**). Strikingly, *Bt* spores remained detectable even after washing, boiling, frying, and microwaving (**Fig. 4B**), highlighting the potential to determine object provenance from cooked foods.

**Fig. 4.**
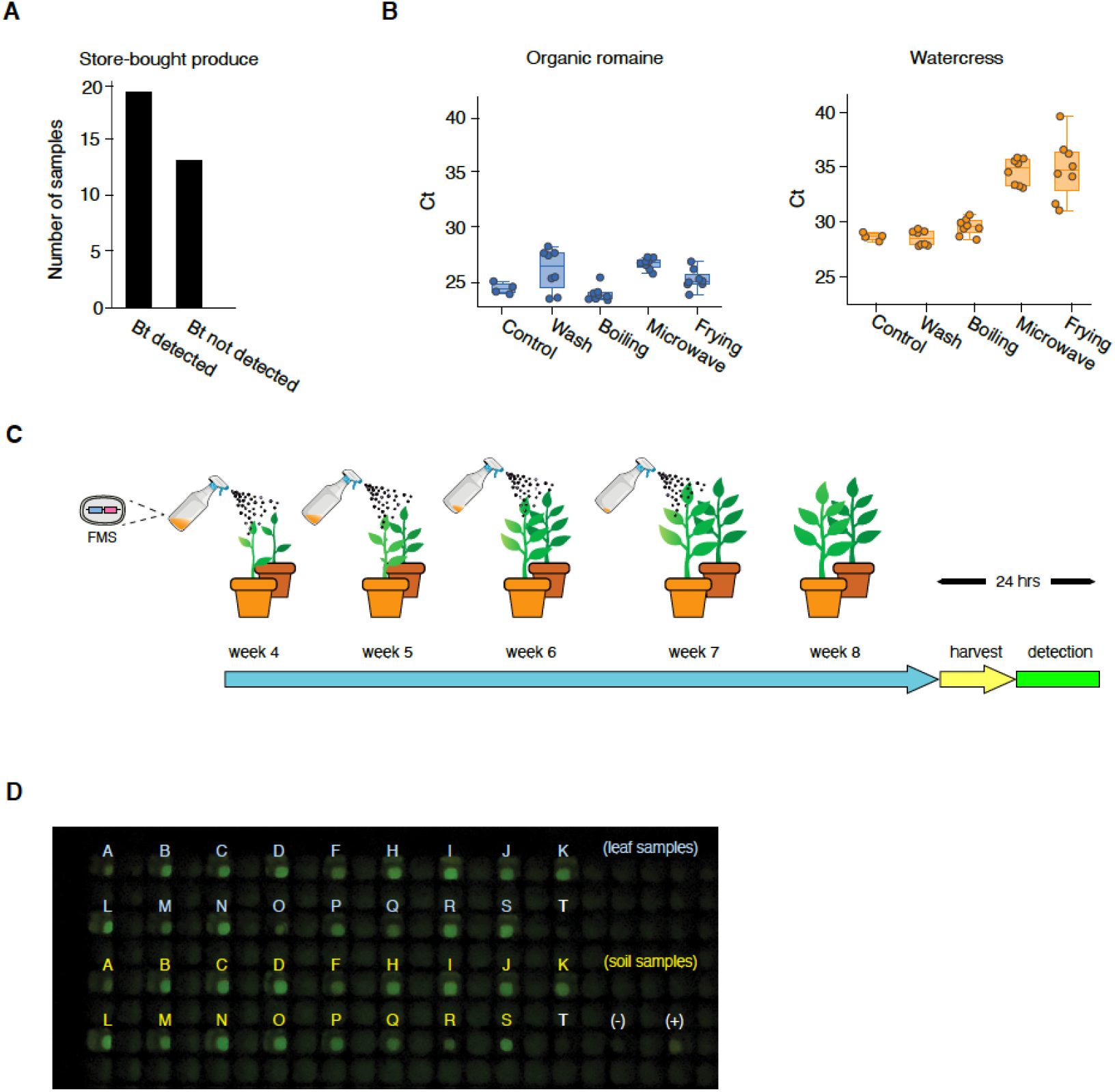
Food provenance of produce using FMS. **(A)** *Bacillus thuringiensis (Bt)* was detected on 19 of 32 store-bought pieces of produce. **(B)** Organic romaine lettuce and watercress, both positive for *Bt*, were washed, boiled, fried, or microwaved. *Bt* remained detectable after these treatments. (n = 2 technical replicates, n = 4 biological replicates for each treatment, n = 2 biological replicates for non-treated control samples). **(C)** 18 distinct *B. subtilis* FMS were applied onto 20 plants in separate pots. FMS were applied once a week (4 times total). **(D)** Detection of FMS on plants and soil after harvesting by SHERLOCK with a group crRNA; mobile phone photograph of reaction/detection plate. Plants A to S were sprayed with FMS, plant T was not sprayed.

Using our barcoded *B. subtilis* FMS as a model for *Bt*, we were able to map FMS on 20 lab-grown leafy plants back to the specific pot in which they were grown. FMS were applied 4 times, beginning 1 week after the first set of leaves appeared, similar to the recommended application protocols for *Bt* spores on crops (**Fig. 4C and fig. S14A**). One week after the final FMS application, the plant and soil from each pot were harvested and tested using SHERLOCK with the group crRNA shared by all but 2 of the FMS. All the samples were positively detected except for the 2 plants with alternate group barcode sequences (**Fig. 4D and figs. S14B-C**). Additionally, using Sanger sequencing, we identified the pot in which each plant was grown for all 19 FMS (**figs. S14, D and E**). The process from DNA extraction to sequence identification took less than 24 hours. This time frame could likely be shortened to hours with massively parallelized hybridization-based detection (*15*).

Importantly, we found that unlike the other surface we sprayed, FMS sprayed on plants did not transfer easily to objects that come in contact with the FMS sprayed plants. The spore transfer efficiency was roughly 1,000-fold less efficient than that of other surfaces such as sand (**fig. S15**). This decrease in transfer efficiency necessitated the use of a more stringent sampling and lysis protocol to remove the FMS from leaves (see Methods). We believe the difference in transfer efficiency is due to FMS adhering to dirt and dust particle; the FMS strongly adhered to these particles as they do to leaves, but the dirt and dust particles can be transferred to other objects. The strong adherence of FMS onto leaves is an advantage for determining food provenance as produce at market would otherwise have signatures of all FMS that the plant came in contact with from harvesting through distribution.

In this work, we showed how rationally engineered microbial spores that can be manufactured in a high-throughput manner, provides a new solution to the object provenance problem. Collectively, our experiments demonstrate that the FMS: 1) persist in the environment, 2) do not spread out of the colonization area, 3) transfer from soil, sand, wood, and carpet to contacting objects and 4) permit sensitive and rapid readout using laboratory and field-deployable methods. The ability to rapidly label objects and determine their trajectories in real-world environments has a broad range of applications across agriculture, commerce, and forensics (*2*). Future iterations of our FMS system could be engineered for limited propagation and actively contained for use in highly trafficked areas, while also allowing for time-resolved information about location history, opening its use to an even wider range of applications.

## Materials and Methods

### Barcode generation

A collection of 28-bp DNA barcodes with a hamming distance greater than 5 were bioinformatically generated. The collection of barcodes was screen against GenBank genome data using BLAST, and any barcodes found to align to genome sequences from a defined isolate were eliminated from the collection.

### Transformation and Barcode Insertion in Bacteria

Bacillus subtilis strains were derived from the wild-type strain 168 and are listed in **Table S2**. Insertion-deletion mutants were from the *Bacillus* knock-out (BKE) collection (*16*). All BKE mutants were back-crossed twice into *B. subtilis* 168 before assaying and prior to antibiotic cassette removal. Antibiotic cassette removal was performed using a temperature-sensitive plasmid encoding the Cre recombinase (*17*).

DNA barcodes were produced by amplifying 164 bp synthetic megaprimers using oligonucleotide primers oCB034 and oCB035 (**Table S3**) in PCR. The barcode fragments were cloned using standard restriction digest cloning into the plasmid pCB018 (*ycgO*::lox66-kan-lox71), a vector for double-crossover integration at the *ycgO* locus.

### Bacterial sporulation

For large scale spore production, *B. subtilis* strains were sporulated in 1 L supplemented Difco Sporulation Medium (DSM) by nutrient exhaustion (*18*) at 37 °C in 4 L flasks. After 36 hours of growth and sporulation, the spores were pellet by centrifugation at 7000 rpm for 30 minutes, washed 2 times with sterile distilled water, incubated at 80 °C for 40 min to kill non-sporulating cells and then washed 5 times with sterile distilled water. Spores were stored at 4 °C in phosphate-buffered saline.

### Evaluating Bacterial Spore Lysis by Microscopy

To rapidly assess the efficacy of different spore lysis protocols, we expressed a fluorescent protein in the spore core and monitored release following lysis by fluorescence microscopy. The *mScarlett* gene was PCR amplified with oligonucleotide primers oCB049 and oCB050 (**Table S3**) from plasmid pHCL147 (*19*), and inserted downstream of the strong sporulation promoter P_*sspB*_ in plasmid pCB137 (*yycR*::*P_sspB_*-spec), a vector for double-crossover integration at the *yycR* locus.

Spores were immobilized on 2% agarose pads. Fluorescence and phase-contrast microscopy was performed using an Olympus BX61 microscope equipped with an UplanF1 100× phase-contrast objective and a CoolSnapHQ digital camera. Exposure time was 400 ms for mScarlett. Images were analyzed and processed using MetaMorph software.

### Transformation and Barcode Insertion in Yeast

Barcodes were introduced into *Saccharomyces cerevisiae* yeast strain BY4743 with standard lithium acetate chemical transformation with a 15min heat shock. Following overnight recovery in YPD media (10 g/L yeast extract, 20 g/L peptone, 20 g/L glucose), cultures were plated on YPD + G418 to select for transformants.

Yeast were transformed with 1ug of barcode oligos, and two linearized plasmids: 50 ng of Cas9 plasmid, F48V (2μ-KanR-pRPL18B-Cas9-tPGK1-GapRepair) and 1 ug of gRNA plasmid, F51V, containing a single gRNA targeting HO and 200-to 300-bp sequences homologous to the GapRepair region in F48V. When both are transformed into yeast cells, the two linearized fragments would assemble into a functional plasmid granting G418 resistance. Once the HO locus targeted by the gRNA was replaced with the barcode sequence, the assembled Cas9 + gRNA plasmid was dispensable (*20*). Plasmids were cured by culturing cells in YPD for overnight followed by spreading cells on YPD plates and replica plated to YPD + G418 plates to select for colonies negative for plasmids.

### Yeast Sporulation

Yeast cells were cultured in 5 mL of YPD medium at 30°C overnight, then transferred to 1 L of YPD medium and cultured for 24 hours. Cells were pelleted by centrifugation at 3000 g for 3 min and washed twice with sterile distilled water. Finally, cells were resuspended in 500 mL of sporulation medium (10 g/L potassium acetate, 1g/L yeast extract, and 0.5 g/L dextrose anhydrate) and incubated at room temperature while shaking for 5 days. Presence of spores was confirmed by microscopy at 60x magnification.

Spores were pelleted and the supernatant was carefully removed. The spores were then washed once then resuspended in 25 mL of sterile distilled water and transferred to 50 mL conical tubes. These tubes were boiled at 100°C for an hour to rupture any remaining vegetative cells. After boiling, spores were pelleted, washed twice then resuspended in 25 mL of distilled water.

### Production of LsCas13a

LsCas13a was purified as described in (*9*), with some modifications. All buffers were made in UltraPure nuclease-free water and all labware used during purification were cleaned with RNase*Zap* before use. Purification of the expressed LsCas13a protein was performed in batch format using StrepTactin sepharose. The SUMO-protease cleaved LsCas13a was concentrated using an Amicon Ultra-0.5 centrifugal filter with a 100 kDa molecular weight cutoff filter. The protein was concentrated until the sample measured as 2mg/mL using the BioRad Protein Assay. The LsCas13a was not concentrated further, and instead stored as 2mg/mL aliquots in lysis buffer supplemented with 1mM DTT and 5% glycerol.

### Recombinase Polymerase Amplification reaction and primer design

Recombinase Polymerase Amplification (RPA) reactions were performed as described in (*9*). The Twist-Dx Basic kit was used according to the manufacturer’s instructions. RPA primers JQ24 and JQ42 (**Table S3**) were used at 480 nM final concentration. Reactions were run with 1 μl of input in 10 μl reactions for 1-2 hrs at 37 °C, unless otherwise noted.

### SHERLOCK detection reactions and crRNAs

Detection reaction reactions were performed as described (*9*). crRNA preparation was performed as described in (*9*), except the *in-vitro* reaction volume scaled to 60 μl. All crRNA and barcode sequences used in this study are available in Tables **S4 and S5**. BioTek readers were used for measuring fluorescence of reactions (Synergy H1 Plate Reader) at Excitation/Emission=485/528 nm wavelength. Positive threshold cutoff value of 2500 was determined by averaging the values of negative control fluorescence values plus 4σ.

### Colonizing surfaces with spores

Spores were diluted to a final concentration ≤1×10^8^ spores/mL in distilled water in order to reduce the viscosity of the solution. Spores were routinely stored at 4 °C for long periods of time, or at room temperature for short periods of time. Diluted spores were sprayed onto surfaces using handheld spray bottles (Fisher scientific). At this concentration, colonization had no visible effect on most surfaces tried, though water stains with white residue did appear on hydrophobically-treated wood due to water beading up on the surfaces.

### Swab collection and lysis

Sterile nylon swabs (Becton Dickinson) were dipped into sterile swab solution (0.15 M NaCl + 0.1% Tween-20) and excess liquid was wiped away. The damp swab was rubbed over the object, covering each part of the surface 2x. The tip of the swab was clipped into a microcentrifuge tube, and 200 uL of freshly prepared 200 mM NaOH was pipetted onto the swab. The tube was heated to 95 °C for 10 minutes, then the base was neutralized with 20 μL 2 M HCl, and buffered with 20 μL 10x TE buffer (Tris-HCl 100 mM, EDTA 10 mM, pH 8.0). Lysate samples were optionally purified with a standard DNA purification AMPure XP bead protocol (Beckman Coulter).

### Spore Quantification by qPCR

Quantitative Polymerase Chain Reactions (qPCR) were prepared in 10μ reactions with SYBR Green I Master Mix (Roche), 1μL of genomic extract as a template, 0.4 mg/mL Bovine Serum Albumin, and 1uM of each primer, listed in **Table S3**. The reactions were carried out in a Roche LightCycler 480 instrument with the following cycling conditions: (i) denaturation, 95°C/10 m (ii) amplification, 45 cycles 95 °C/10 s, 60°C/5 s, 72°C/10 s.

### Design of ~1 m^2^-scale test surfaces and perturbations

We constructed small-scale test surfaces in an incubator to simulate real world conditions in which the Forensic Microbial Spores may be deployed. We assembled 20 test surfaces, from 4 materials (sand, soil, carpet, and wood), divided into control and perturbed conditions (**Table S7**). Each surface was divided into a 9” × 13” grid, denoting different locations for direct samples and transfer samples each week. The gridded area of each surface was colonized with different pairs of barcoded strains, using a handheld spray bottles, to a final concentration of roughly 7.5 × 10^6^ *B. subtilis* spores and 2.5 × 10^5^ *S. cerevisiae* spores per square inch.

For outdoor conditions, twelve 9” × 13” trays were filled to a 1” depth with either sand or potting mix, and housed on shelved in one of two incubators (Shel Labs) heated to 25 °C (**fig. S4A**). To simulate wind, one incubator was equipped with 140 mm computer fans, one directed at each of the 6 perturbed trays. Simulated rain was applied to perturbed surfaces each week, varying in intensity from handheld spray bottle (200 mL / week) on weeks 1-6 to watering can (500mL / week) on weeks 7-12. The incubator housing the perturbed surfaces was also programmed to fluctuate in temperature between 25 °C and 35 °C with a period of one week.

For indoor conditions, we cut out 4 sections of carpet and 4 sections of laminate wood flooring and marked out 9”×13” sections on each for testing. All eight surfaces were shelved in an incubator (Shel labs) heated to 25 °C, and humidified to 40-50% RH (**fig. S4**). To simulate cleaning, perturbed carpet surfaces were cleaned using a handheld vacuum, while perturbed wood surfaces were cleaned with a hand broom (**fig. S4A**).

### Sampling from ~1 m^2^-scale test surfaces over a three-month period

Each week for a 13-week period, 0.25 g of sand or soil was sampled from each surface and processed using a DNeasy Powersoil kit (Qiagen) to isolate DNA, following the manufacturer’s protocol. For carpet and wood samples, we performed our swab collection and lysis protocol to generate lysate for barcode qPCR. For all surfaces, 1” × 2” test objects (either rubber or plywood) were used to test transferability of spores; test objects were pressed onto the surface a single time, then processed with the swab collection and lysis protocol to generate lysate used for barcode qPCR without AMPure XP bead cleanup.

### Design of full-scale sandpit and perturbations

A 6 m × 16 m × 0.25 m indoor sandpit was built and equipped with drainage and a slight grade. A 1 × 6 m section along the top edge was colonized with *B. subtilis* BC-24 & BC-25 spores with roughly 2.5 × 10^11^ spores each, and *S. cerevisiae* BC-49 & BC-50 spores with roughly 1.25 × 10^10^ spores each. One half of the sand pit was designated for environmental perturbations, with 1 m diameter fans placed at the colonization end, and hose simulated rain (0.5 in / week) applied each week.

### Sampling from full-scale sandpit over a three-month period

Each week for a 13-week period, ~0.2 g of sand or soil was sampled from 20 locations in the sandpit and processed using the NaOH lysis protocol. For testing transferability, objects (shoes or wood) were pressed onto the surface a single time (**fig. S7**), then processed with our sample swab and spore lysis protocol. In this experiment, lysates were used for SHERLOCK detection without prior AMPure XP bead cleanup.

### Measuring spore retainment on shoes

24 pairs of shoes were worn in the colonization zone of the 64 m^2^ sand pit, accumulating spores over a 1-minute walking period. These shoes were then divided into 8 groups, and were worn while walking in uncolonized areas for 0, 1, 5, 15, 30, 60, 120 or 240 minutes. Shoes were processed with the swab collection and lysis protocol, followed by AMPure XP bead purification. Purified DNA was used as a template for SHERLOCK detection and qPCR quantification.

### Determining Object Trajectory

Grids of varying sizes (averaging 0.25 m^2^ per square) and materials (sand, soil, carpet, wood) were colonized using approximately 2.5 × 10^11^ *B. subtilis* spores and/or 1.25 × 10^10^ *S. cerevisiae* spores per square. Shoes and remote-control cars were used as test objects, and were exposed to colonized surfaces by walking or driving on the test grid surfaces. All surfaces of test objects were swabbed and processed with our spore lysis protocol. For 1 unique FMS / sand quadrant, a quadrant is called positive if SHERLOCK reaction is positive for the unique FMS of that quadrant; for 2 unique FMS / sand quadrant, a quadrant is called positive if SHERLOCK reaction is positive for at least 1 of the 2 FMS of that quadrant; for 4 unique FMS/sand quadrant, a quadrant is called positive if SHERLOCK reaction is positive for at least 2 FMS of that quadrant.

### Qualitative Readout of SHERLOCK using mobile phone camera

To mimic in-field deployment, a portable light source, an orange acrylic filter was set up for data collection (**fig. S10B**), a mobile phone at default setting (flash off) collected signals after SHERLOCK reactions had proceeded 30-60 minutes in dark box or dark room.

### PCR of *Bacillus thuringiensis* on produce

Around 250 mg of each produce sample was cut into 1-3 mm pieces using a scalpel and then processed with the DNeasy Powersoil Pro kit (Qiagen) to isolate 50μL of eluted DNA, following the manufacturer’s protocol. Then, 1 μl of the DNA was subjected to PCR with primers BT-1F and BT-1R (**Table S3**) using Kapa Biosystems HiFi HotStart ReadyMix (Thermofisher Scientific) with the following cycling conditions: (i) denaturation, 95°C/3 m (ii) amplification, 40 cycles 98°C/20 s, 60°C/15 s, 72°C/30 s.

### Robustness of *Bacillus thuringiensis* on produce

We selected produce that was positive for *Bacillus thuringiensis*, then treated these samples with various cooking methods: washing, boiling, microwaving, or frying. For washing, the produce pieces were placed in a 50 ml conical tube covered with a screen with tap water running over the sample for 10 min then dried in paper towels. For boiling, a piece of produce was placed in an Eppendorf tube filled with 1ml of water and placed in a boiling beaker of water for 15 min then dried in paper towels. For microwaving, produce pieces were placed in a petri dish with the cover on at full power for 2 min. For frying, 1ml of vegetable oil was added to a 250 ml or 400 ml beaker that was pre-heated for 1min on hotplate set to 350 °C then the piece of produce was added and heated for 1 min with occasional stirring. Around 250 mg of cooked sample was processed with DNeasy PowerSoil Pro Kit (QIAGEN). qPCR was performed using primers BT-1F and BT-1R (**Table S3**) and PowerUp SYBR Green Master Mix (ThermoFisher Scientific).

### Barcode identification from a model farm

Twenty garden pots were filled with potting mix and enclosed with canvas with 12 hours of daily blue light. One seedling was planted in each pot. Temperature was controlled to be around 23 °C. Plants were watered every 2-3 days and exposed to blue light for 12 hours daily. Barcoded *B. subtilis* spores were sprayed on the plants separately once a week for 4 weeks during the growth period. In total, ~10^8^-10^9^ spores were sprayed on each plant. One week after the final spraying, plant samples were harvested and processed using DNeasy PowerSoil Pro Kit as described above to isolate DNA. Barcode DNA was amplified using BTv2-F and BCv2-R (**Table S3**) using Kapa Biosystems HiFi HotStart ReadyMix, then Sanger sequencing was used to identify the barcode sequences (GENEWIZ).

## Acknowledgments

We thank participating labs’ members for useful feedback; Ofer Mazor the HMS Research Instrumentation Core Facility for technical consultations and instrument design and fabrication. This facility is generously supported by the HMS Department of Neurobiology, the Bertarelli Program in Translational Neuroscience and Neuroengineering at HMS, and by an NEI P30 Core Grant for Vision Research #EY012196.

## Author contributions

JQ, ZL, CPM, HYJ, RCB-O, SB conceived the study, performed most key experiments and data analysis (supervised by MS). JQ, ZL, CPM, HYJ, RCB-O, SB and FHR-G wrote the paper (with MS, DR). VJ, AS, KS, LA, GJ, MA, MM, LL, SVO assisted with technical experiments. SB purified Cas13 proteins used in this study. RCB-O and FHR-G designed and constructed *B. subtilis* strains and purified spores (supervised by DR). DR, PS, MB provided critical insights used in this study. All authors reviewed the manuscript.

## Data and materials availability

All data are available in the article or the supplementary materials.

## Supplementary Materials

Materials and Methods

Figures S1-S15

Tables S1-S7

