## Supplementary Materials for "Forensic microbial system for high-resolution object provenance"

**This PDF file includes:**

Supplementary Text
Figs. S1 to S15
Tables S1 to S7

**Supplementary Text**

Developing high-throughput methods to screen barcodes and crRNAs

In order to scale the system, we devised a facile method to quickly screen a large number of barcodes and crRNAs in parallel to eliminate those with high cross reactivity or background; we performed pooled *n-1 barcodes* RPA reactions with a corresponding crRNA and water RPA controls. A pooled *n-1 barcodes* RPA reaction denotes a single RPA reaction with all DNA barcodes pooled together except for barcode that is left out, and this pooled reaction is screen against all crRNAs. Following our original in-vitro screen of 22 barcode crRNA pairs, we made 72 additional barcodes and tested all 94. Retesting our initial 22 crRNA validated our assay; 19 of the 22 passed while crRNAs 7, 11, and 12 that were just below our cut-off in our pairwise test (see Methods) scored as cross-reactive in the pooled assay. In total, we eliminated 17 of the 94 crRNAs due to high background and 7 additional crRNAs were eliminated due to cross-reactivity with multiple barcodes. Cross-reactivity appeared to be caused by crRNAs instead of barcodes as no barcodes were cross-reactive with all guides (**figs. S1, B and C**).

Feasibility testing of Forensic Microbial Spores on incubator-scale test surfaces

Compared to in-vitro tests, real-world environments often present challenges for enzyme-based analyte detection systems, both by sequestering analyte and inhibiting reactions. Real-world environments also present a challenge to the stability of forensic labels, by degrading or washing away the label over time. We tested the feasibility of our forensic microbial spores by constructing chambers to simulate different real-world environments and perturbations. We chose four material types: sand, soil, carpet, and wood, and housed test surfaces of these materials in modified incubators in order to simulate indoor and outdoor conditions. We also devised perturbations (e.g. rain/wind/cleaning) that might be expected to remove spores from the surface.

Over a three-month period, qPCR targeted to our forensic microbial spores demonstrated no significant loss of spores over time (**Fig. 2A**). Importantly, the perturbations did not significantly reduce detection compared to control surfaces for any of the 4 materials. Furthermore, in most cases, spores could be transferred to a rubber or wood test object after a single direct exposure to the colonized surface, and subsequently detected by qPCR (**fig. S4**).

Sensitivity and specificity of PCR-based detection of *Bacillus thuringiensis*

To understand the sensitivity of our primers, we used qPCR (described below) to test different amounts of purified *Bt* gDNA, where there should be no inhibition from the spores. The method is sensitive enough to detect the amount of gDNA equivalent to as few as less than 10 spores (**fig. S13A)**. Furthermore, we tested 14 produce samples from local farms that we know had sprayed *Bt* on the produce, and *Bt* was detected in all of them (**fig. S12**).

To test the specificity, we included 4 produce samples from personal gardens that had not been sprayed with *Bt*, and two samples from our lab farm, which also had not been sprayed with *Bt*. We also included 5 unrelated plants from 5 different places. Those 10 samples served as negative controls. We did not detect any signal from them (Samples 10 to 13 & 17 in **fig. S13B** and Samples 1 to 5 in **fig. S12B**). To further test the validity of the detected bands from produce, we randomly selected 6 produce **(fig. S13B** sample 1-6**)** and send their PCR products to Sanger sequencing. The sequences matched the cry1A genes and belonged to three variants (**fig. S13C**). When the sequences were tested with NCBI BLAST against the nr database, the only organism showed up in the top 100 hits was *Bt* (not shown). Lastly, we tested a pool of non-*Bt* bacterial gDNA (*Streptomyces Hygroscopicus*, *S. cerevisiae*, *B. subtilis*, *E. coli* and *Pseudomonas*) along with *Bt* gDNA and only *Bt* gave rise to PCR bands **(fig. S12C).**

Figure. S1

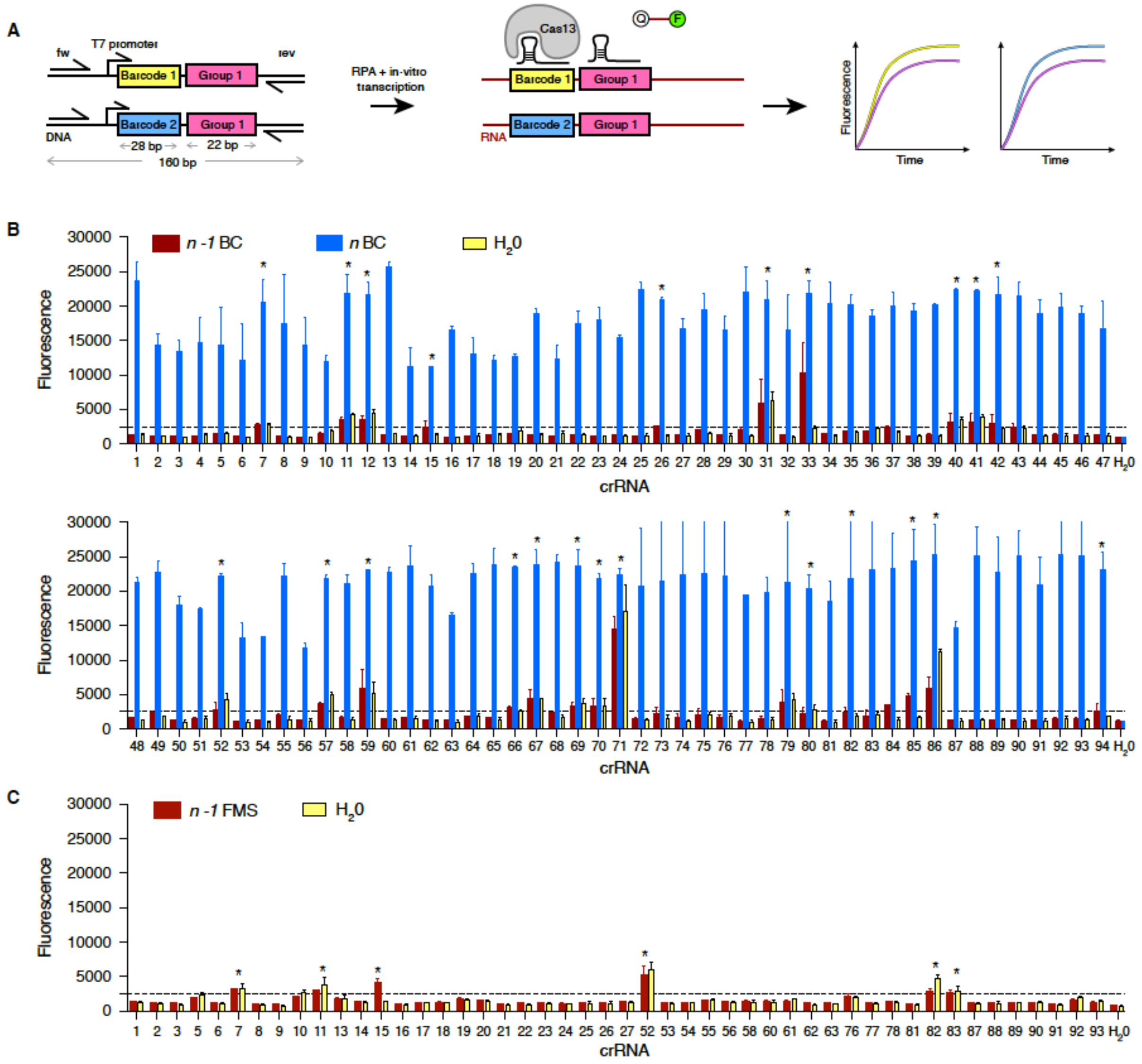


**Fig. S1. Screening for cross-reactive crRNA-barcode pairs.** (**A**) Schematic of FMS detection pipeline SHERLOCK. Schematic of DNA barcode region design (160 bp) with RPA primers. Specific barcodes (28 bp) are highlighted in yellow and blue. Group barcode (22 bp) is highlighted in pink. (**B**) *in vitro* DNA barcode and crRNA cross-reactivity assay. Bars depict SHERLOCK signal from reactions using different templates, with the *n-1* barcodes reactions in red, all barcodes reactions in blue, and water RPA reactions in yellow. * denotes crRNA with high background or cross-reactivity. (n = 3 technical replicates, error bars represent mean + s.e.m.). (**C**) *in vivo* barcoded-FMS and crRNA cross-reactivity assay. Shown are the *n-1* FMS reactions in red and water RPA reactions in yellow. * denotes crRNA with high background or cross-reactivity (n = 3 technical replicates, bars represent mean + s.e.m.).

Figure. S2

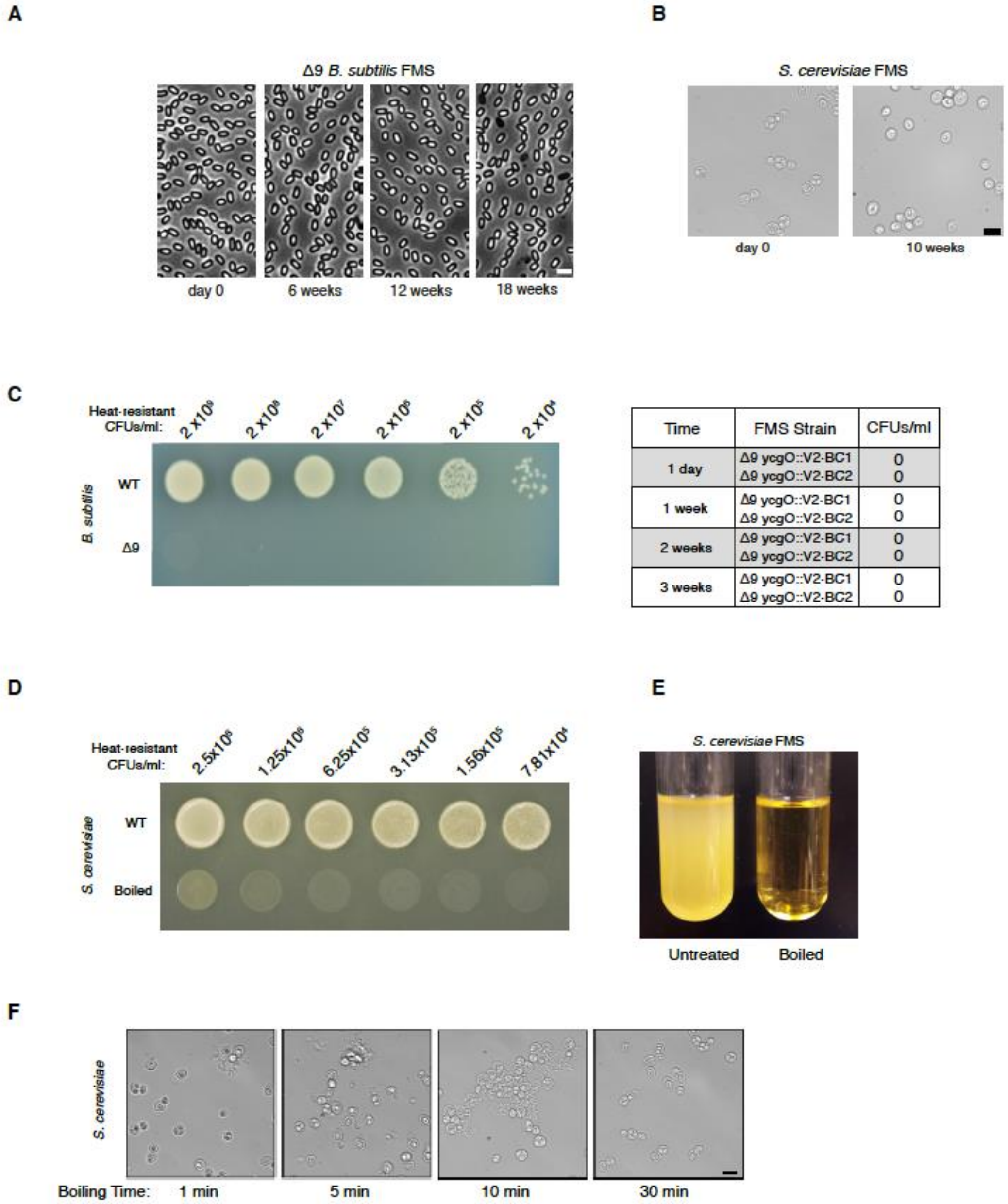


**Fig. S2. Germination defective FMS are biocontained. (A**) *B. subtilis* Δ9 FMS remain stable and dormant over 4 months when stored in PBS at room temperature. At the indicated time points, spores were analyzed by phase-contrast microscopy. Scale bar indicates 2µm. **(B)** *S. cerevisiae*

FMS remain stable after 10 weeks. Images of FMS at day 0 and week 10 are shown. Asci remained intact and no cell lysis was observed. Scale bar indicates 10 µm. (**C**) *B. subtilis* Δ9 FMS are unable to germinate, outgrow, and form colonies on nutrient rich medium. ~2 x $10^9$ WT and Δ9 FMS were 10-fold serially diluted in PBS and 10µL spotted on LB agar. Plates were incubated at 37 ˚C for 16 h. At the indicated time points, ~2x$10^9$ Δ9 FMS with the indicated barcodes (BC-1 and BC-2) were plated on LB agar. After 16 h at 37˚C the number of colony-forming units were counted. (**D**) *S. cerevisiae* FMS are unable to germinate, outgrow, and form colonies on nutrient rich medium after boiling. 2.5x$10^6$ cells before and after 1 hour of boiling were either serially diluted at 2-fold and spotted on YPD. Note that the dim circles in the boiled sample on the YPD plate are cell debris and do not indicate growth. **(E)** *S. cerevisiae* Cultured in liquid YPD and incubated at 30˚C overnight. Untreated cells showed substantial growth while boiled FMS showed no growth. **(F)** Images of the effect of boiling on *S. cerevisiae* vegetative cells and FMS. Even after 1 min, the majority of the intact cells are FMS. At 30 min, all the intact cells observed are FMS. Scale bar indicates 10 µm.

Figure. S3

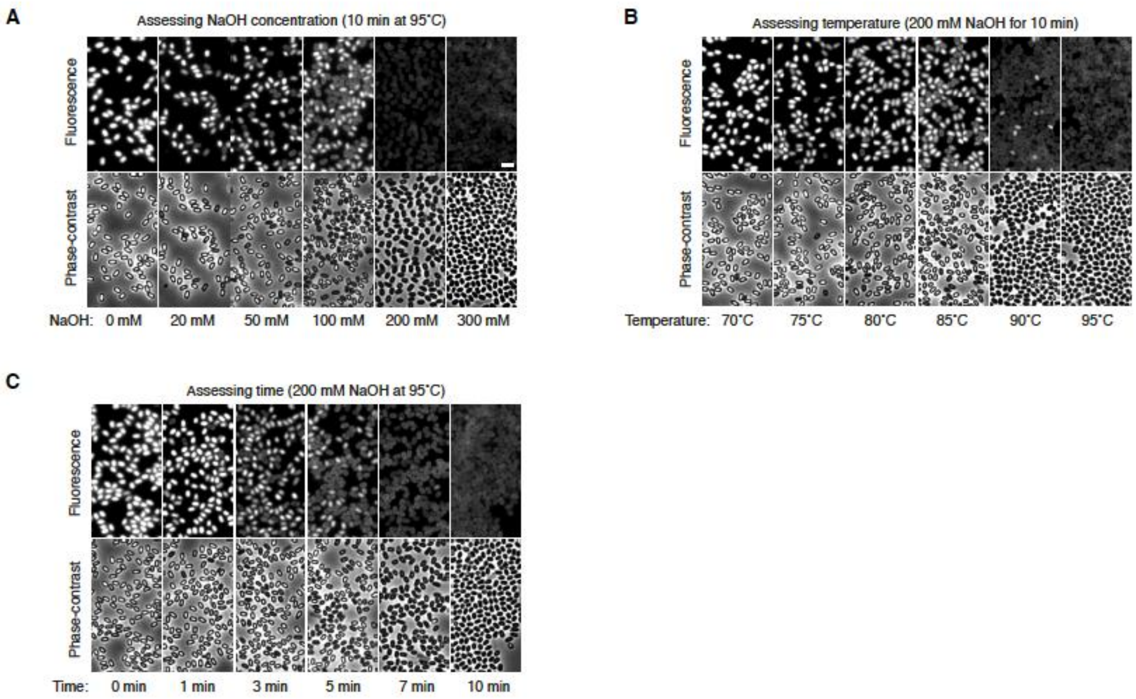


**Fig. S3. Optimization of *B. subtilis* FMS lysis.** (**A**) To rapidly assess the efficacy of lysis, Δ9 FMS harboring a cytoplasmic red fluorescent protein (mScarlet) were analyzed by fluorescence and phase-contrast microscopy after treatment. ~2 x$10^6$ Δ9 FMS were resuspended in 50 µL of NaOH at the indicated concentrations and heated for 10 min at 95 ˚C. (**B**) ~2 x$10^6$ Δ9 FMS were resuspended in 50 µL of 200 mM NaOH and heated at the indicated temperatures for 10 min. (**C**) ~2 x$10^6$ Δ9 FMS were resuspended in 50 µL of 200 mM NaOH and heated at 95˚C for the indicated amount of time. After treatment, FMS were pelleted, washed and resuspended in PBS. An aliquot was then analyzed by fluorescence and phase-contrast microscopy. Loss of fluorescence correlated with the transition from phase-bright to phase-dark FMS. Scale bar indicates 2µm.

Figure. S4

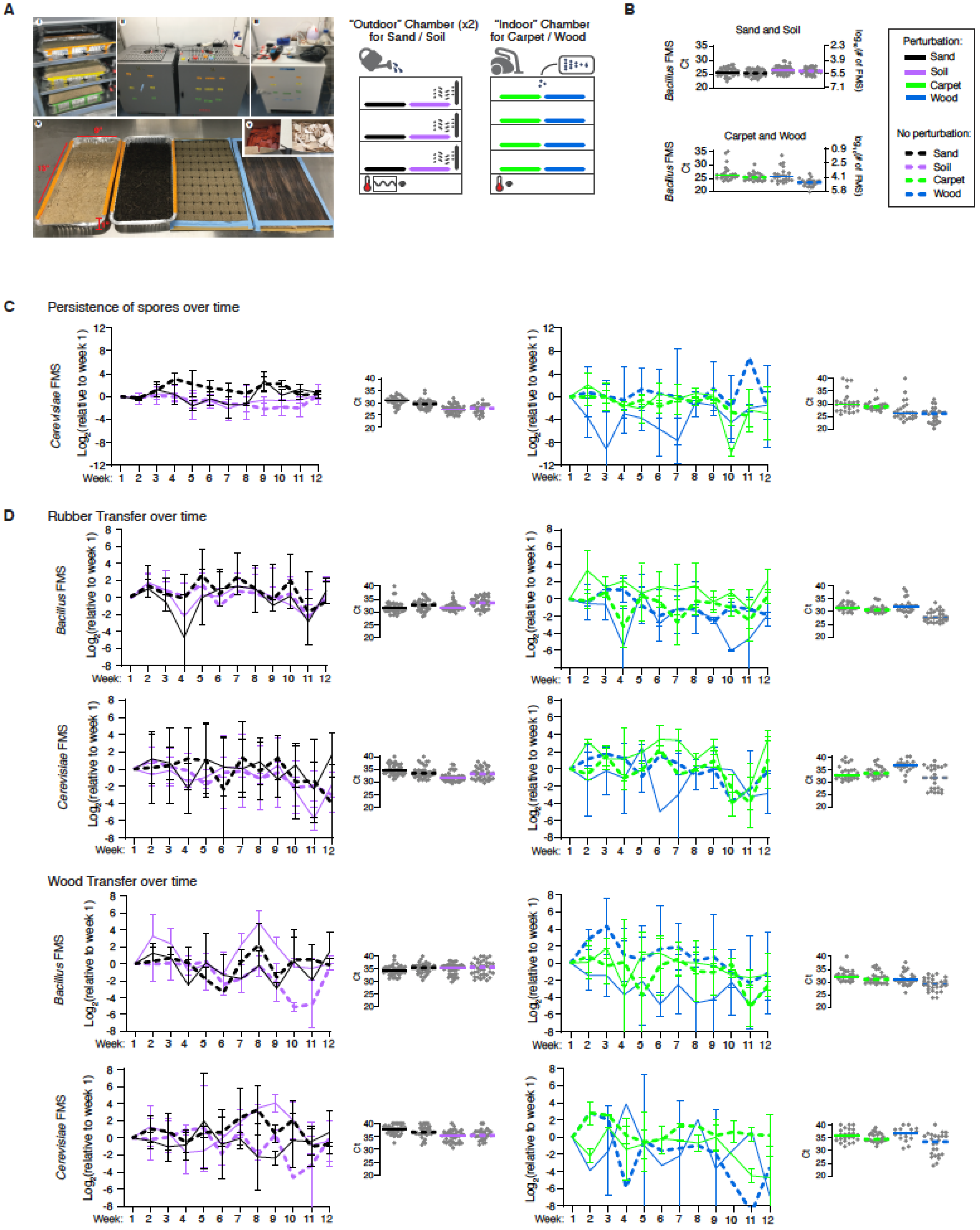


**Fig. S4. Persistence, transferability and maintenance of FMS.** (**A**) Photos and schematic of lab incubator scale experiments and simulated wind, rain, and vacuuming. **(B)** Dot plot of real-time qPCR Ct values (left y-axis) and FMS numbers (right y-axis) based on a qPCR standard curve (in **fig. S8**). (**C**) FMS persisted on sand, soil, carpet and wood surfaces for at least 3 months. FMS count number (relative to week 1 values) and qPCR Ct values. (**D)** FMS were transferable for at least 3 months after colonization from all 4 test surfaces. Rubber and wood objects placed on colonized surfaces were used for testing FMS transferability. Relative FMS count numbers and qPCR Ct values.

Figure. S5

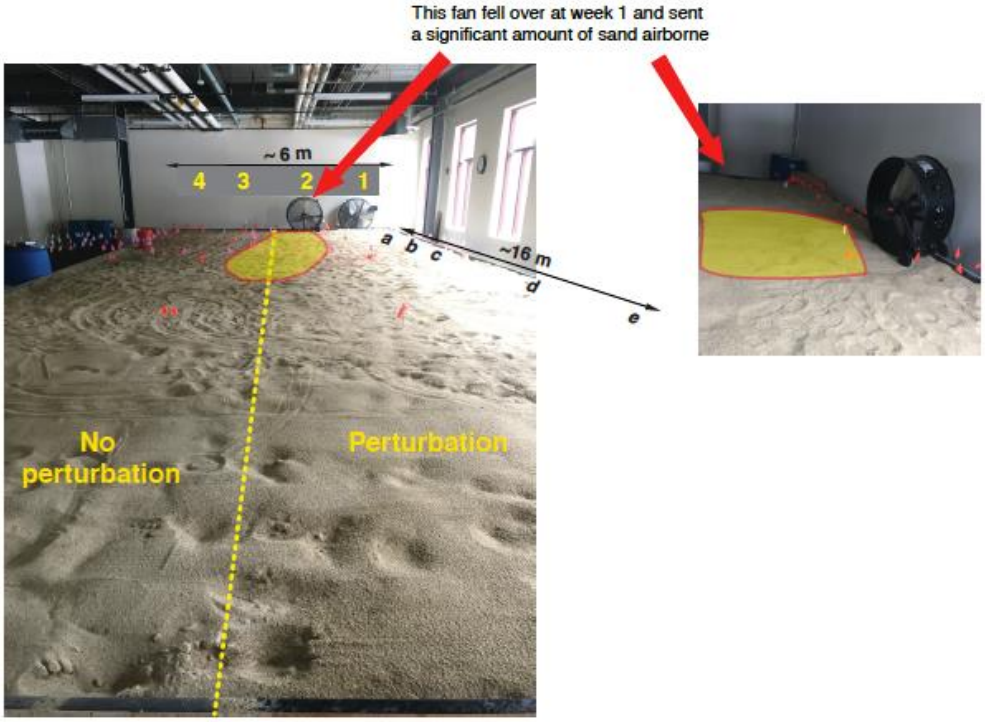


**Fig. S5. Catastrophic disturbance.** Photographs of the large-scale sandpit, with the disturbance area from the fan falling over indicated in yellow.

Figure. S6

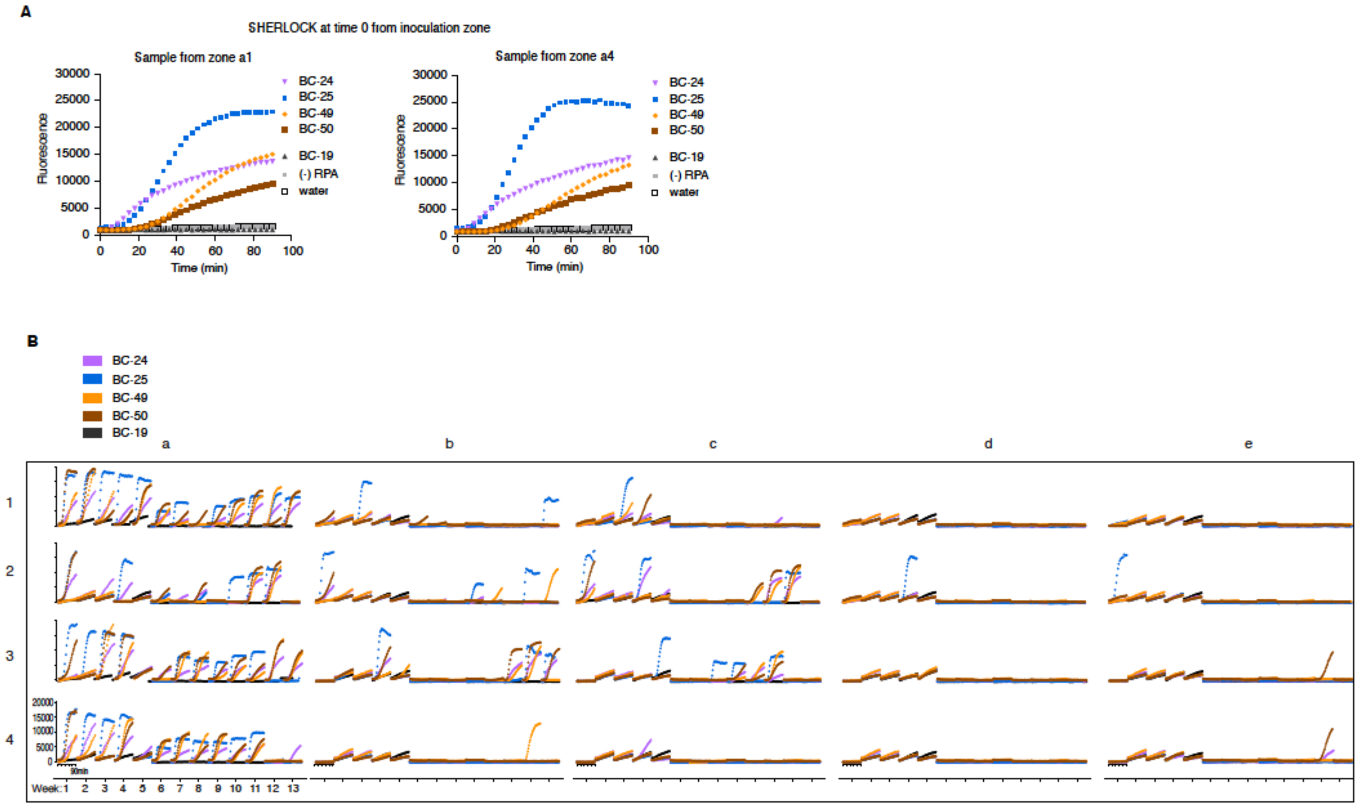


**Fig. S6. Persistence of FMS over time.** (**A**) SHERLOCK detection of BC-24, 25, 49, 50 FMS from samples taken immediately after FMS inoculation, shown as fluorescence timecourses. BC-19, water, and (-) RPA are negative controls. **(B)** SHERLOCK detection indicates FMS persistence over 3 months with or without perturbation. SHERLOCK timecourse data (y-axis: fluorescence, x-axis: time in minutes) is shown for each barcode (colors) at each of 20 locations (a-e, 1-4) for 13 weeks. Signal above the threshold are counted and scored in Fig 2D.

Figure. S7

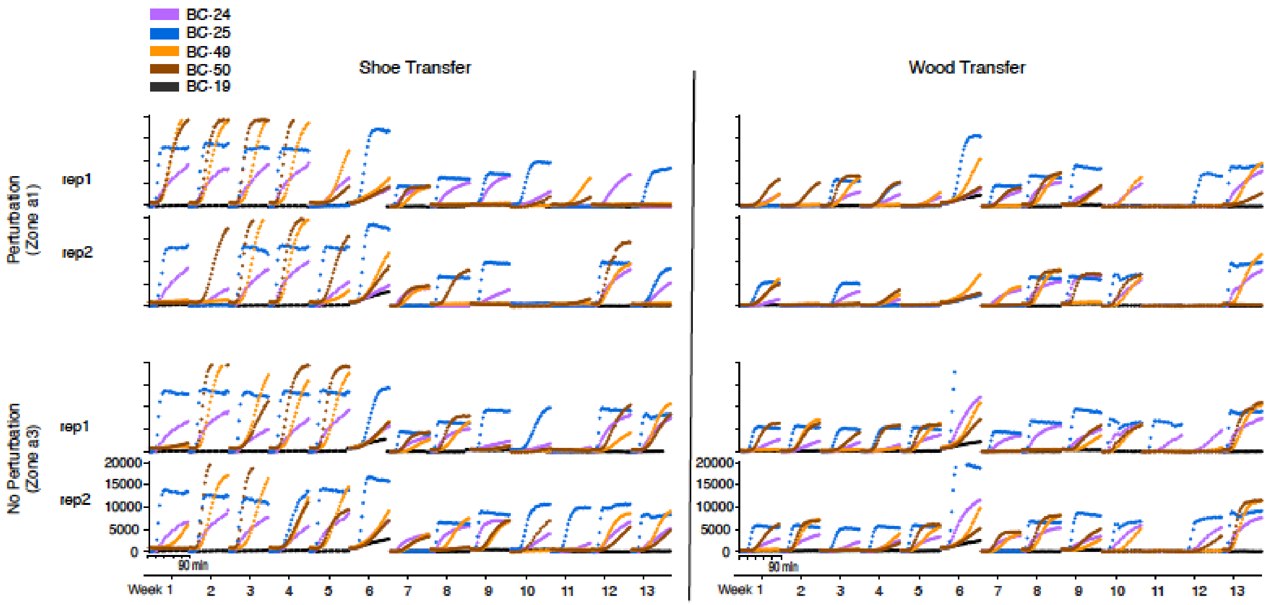


**Fig. S7. Transfer of FMS over time.** SHERLOCK can detect FMS on shoes and wood that come in contact with a colonized surface. SHERLOCK timecourse data (y-axis: fluorescence, x-axis: time in minutes) for transfer onto shoes or wood is shown for each barcode (colors), for 2 replicates from 2 different colonized locations (a1 and a3) over 13 weeks. BC-19 is a negative control.

Figure. S8

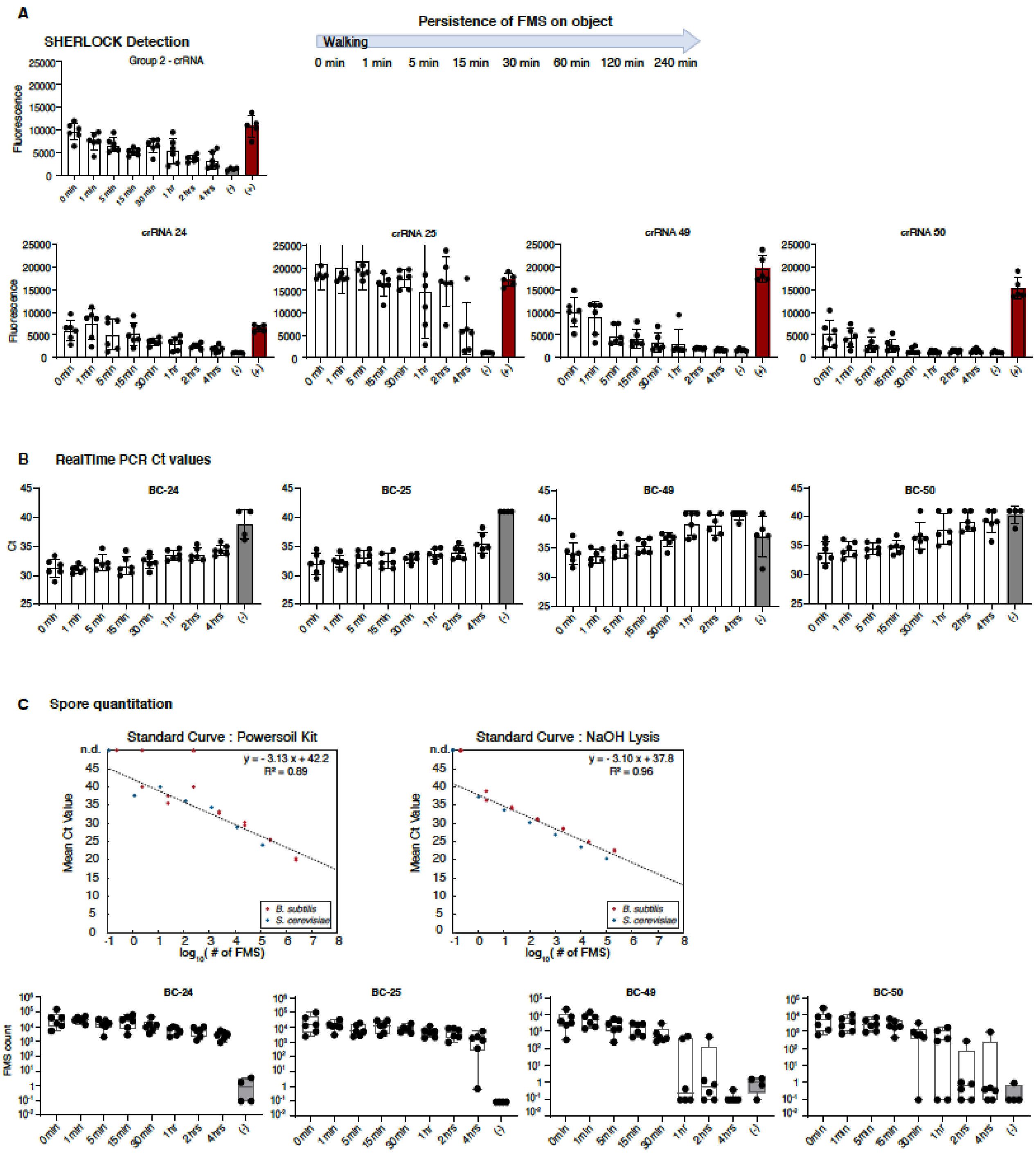


**Fig. S8. Persistence of FMS on object after transfer.** (**A**) SHERLOCK and (**B**) qPCR showing BC-24 and 25 *B. subtilis* FMS and BC-49 and 50 *S. cerevisiae* FMS retained on the shoe after up to 4 hours of walking on uncolonized areas. (**C**) Standard curves constructed from known FMS quantities using either PowerSoil or NaOH lysis techniques. (**D**) Estimating the number of FMS on each shoe from Ct values using the qPCR standard curve.

Figure. S9

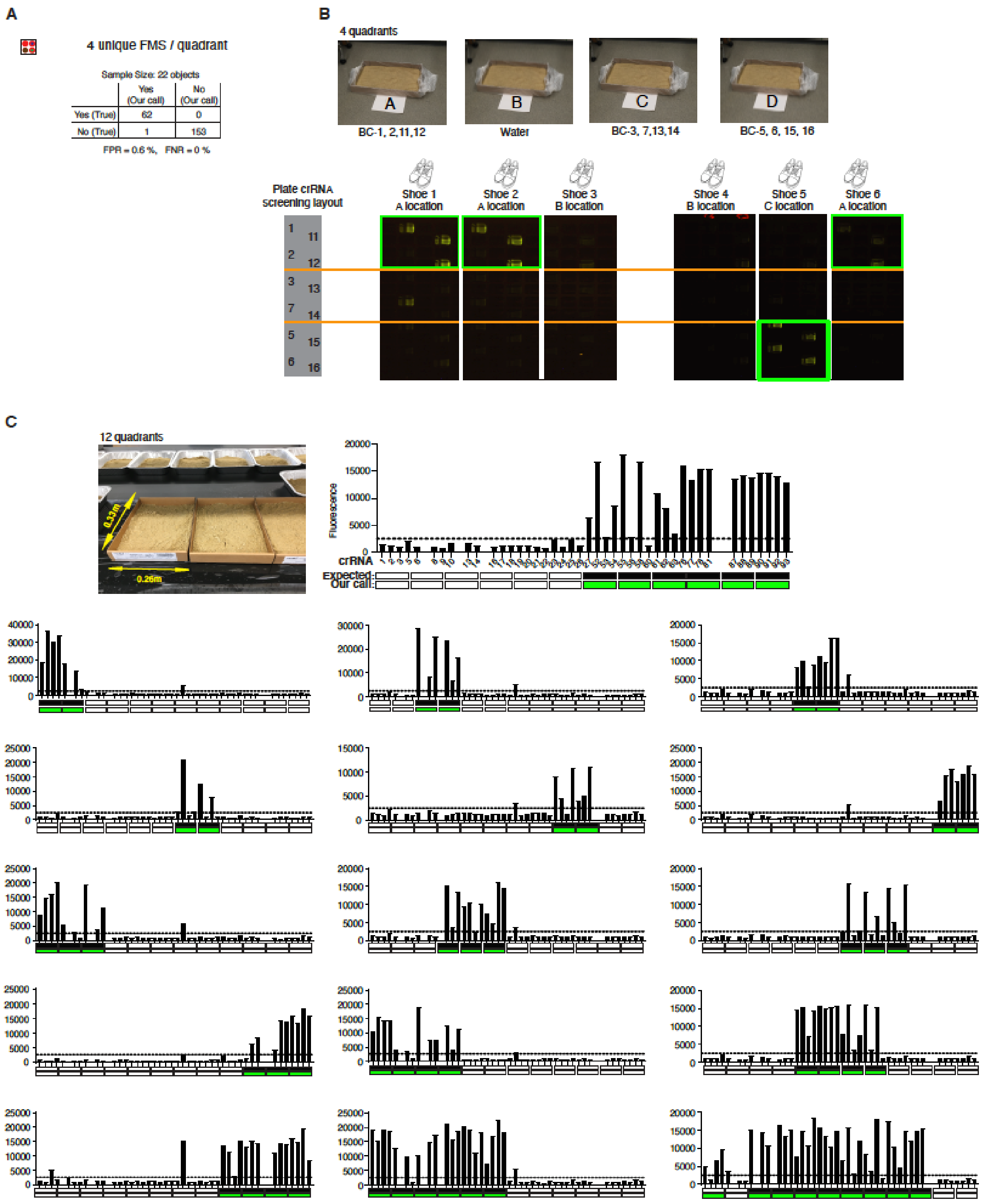


**Fig. S9. Object provenance using 4 unique FMS per quadrant. (A)** see **Fig. 3C**. (**B**) 6 shoe samples were tested in 4 quadrants labelled with 4 FMS per quadrant (photograph of representative quadrant) and SHERLOCK reaction plates imaged using a mobile phone. (**C**) Another 16 samples were tested in 12 similar quadrants; bar plots depict endpoint SHERLOCK signals.

Figure. S10

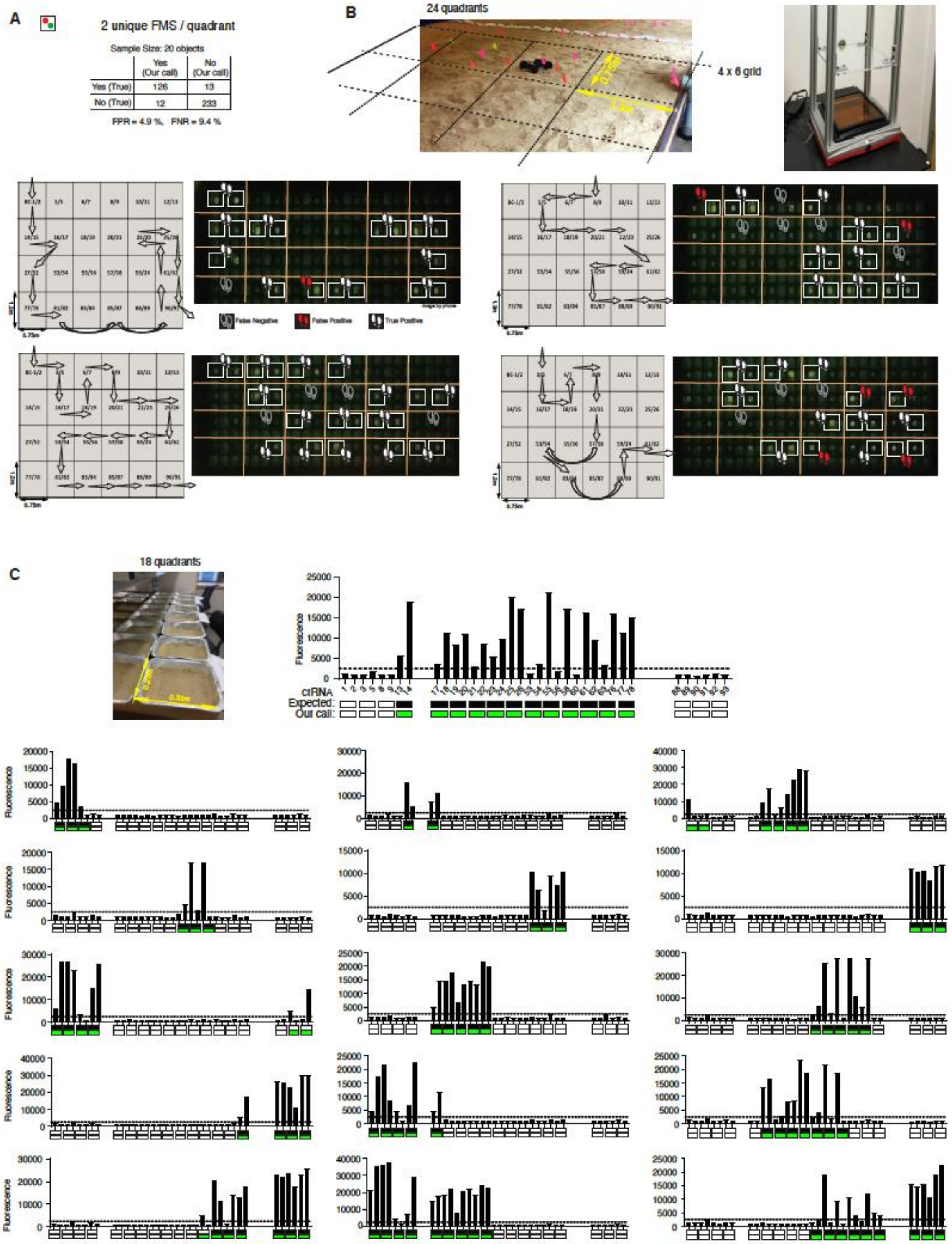


**Fig. S10. Object provenance using 2 unique FMS per quadrant.** (**A**) False positive and false negative rates for barcode detection with 2 unique FMS per quadrant. (**B**) (**Left**) 4 shoe samples were tested in 24 quadrants labelled with 2 FMS per quadrant (photograph of representative quadrant) and SHERLOCK reaction plates were imaged using a mobile phone. (**Right**) Photograph of the apparatus for taking plate images in the field. (**Bottom**) In the diagram to the left of each image, arrows indicate the real walking path; shoe prints in image indicates our prediction. (**C**) 16 samples tested in small-scale with a sand surface, were tested by SHERLOCK as above.

Figure. S11

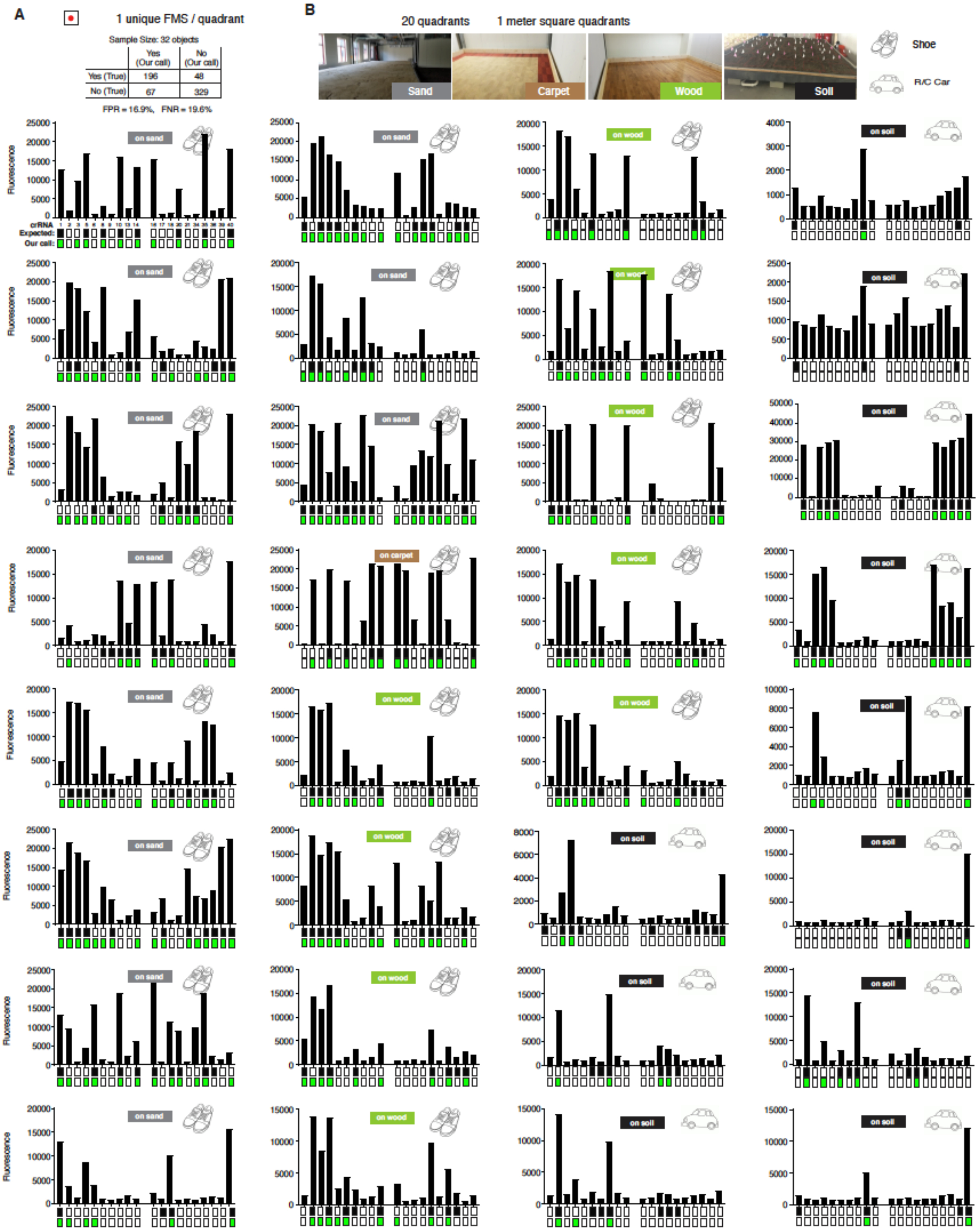


**Fig. S11. Object provenance using 1 unique FMS per quadrant. (A)** False positive and false negative rates for barcode detection with 1 unique FMS per quadrant (**B**) 32 samples walked on different surfaces were tested by SHERLOCK as described in **fig. S10**.

Figure S12

A

| Test Result | Positive Control | Negative Control | Total |
|---|---|---|---|
| Positive | 14 | 0 | 14 |
| Negative | 0 | 10 | 10 |
| Total | 14 | 10 | 24 |

B

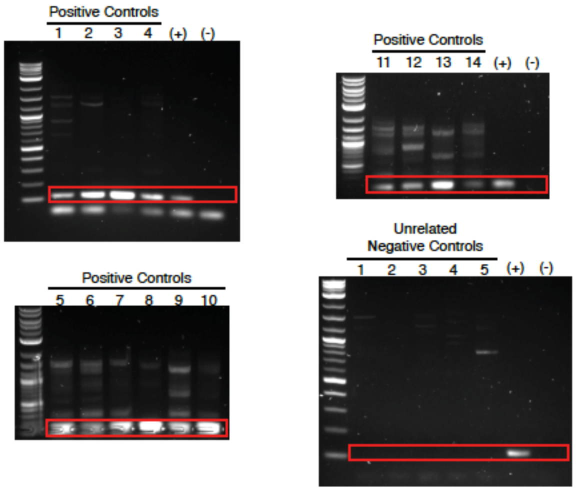


C

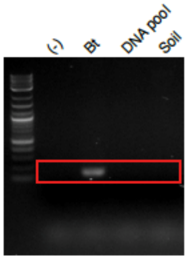


**Fig. S12. Detection of *Bacillus thuringiensis* controls. (A)** Positive controls of *Bt* (i.e. produce sprayed with *Bt* during growth) all tested positive and negative controls (i.e. plants from personal gardens or other plants known not to be sprayed with *Bt*) all tested negative by PCR. **(B)** PCR gel images of *Bt* positive and negative control samples from (**A**). Red box indicates the *Bt* Cry1A band. *Bt* gDNA was used as a positive control template (+); water was used as negative control template (-). **(C)**. Specificity of PCR-based *Bt* detection. Different template samples including *Bt* gDNA, a DNA pool of gDNA from non-*Bt* microbes (*Streptomyces hygroscopicus*, *S. cerevisiae*, *B. subtilis*, *E. coli* and *Pseudomonas*), a soil sample (soil) or water (-) were subjected to PCR-based *Bt* detection.

Figure. S13

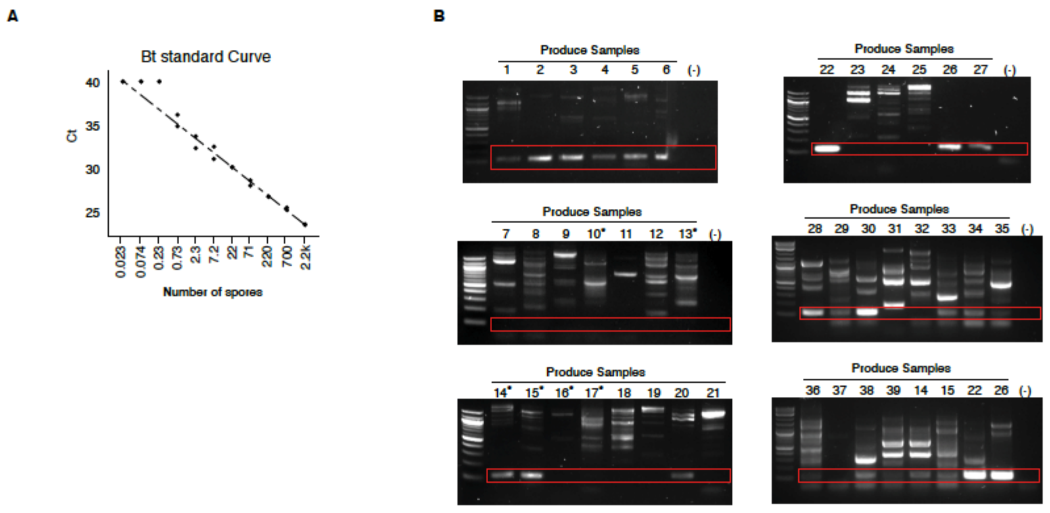


C

```
                       2001                                   2040
cry1A                  GTCCGAGAAAGTCAAACATGCGAAGCGACTTAGTGATGAG
organic_kale.........  ------------CAAACATGCGAAGCGACTTAGTGATGAG
organic_spinach......  ------------CAAACATGCGAAGCGACTTAGTGATGAG
organic_lettuce......  ------------CAAACATGCGAAGCGACTTAGTGATGAG
non-organic_kale.....  ----------------CATGCGAAGCGACTTAGTGATGAG
non-organic_spinac...  -----------TCAAACATGCGAAGCGACTTAGTGATGAG
non-organic_lettuc...  ---------------ACATGCGAAGCGACTTAGTGATGAG

                       2041                                   2080
cry1A                  CGGAATTTACTTCAAGATCCAAACTTTAGAGGGATCAATA
organic_kale.........  CGGAATTTACTTCAAGATCCAAACTCTAGAGGGATCAATA
organic_spinach......  CGGAATTTACTTCAAGATCCAAACTTTAGAGGGATCAATA
organic_lettuce......  CGGAATTTACTTCAAGATCCAAACTTTAGAGGGATCAATA
non-organic_kale.....  CGGAATTTACTTCAAGATCCAAACTCTAGAGGGATCAATA
non-organic_spinac...  CGGAATTTACTTCAAGATCCAAACTTCAGAGGGATCAATA
non-organic_lettuc...  CGGAATTTACTTCAAGATCCAAACTTCAGAGGGGTCAATA

                       2081                                   2120
cry1A                  GACAACTAGACCGTGGCTGGAGAGGAAGTACGGATATTAC
organic_kale.........  GACAACTAGACCGTGGCTGGAGAGGAAGTACGGATATTAC
organic_spinach......  GACAACTAGACCGTGGCTGGAGAGGAAGTACGGATATTAC
organic_lettuce......  GACAACTAGACCGTGGCTGGAGAGGAAGTACGGATATTAC
non-organic_kale.....  GACAACTAGACCGTGGCTGGAGAGGAAGTACGGATATTAC
non-organic_spinac...  GACAACTAGACCGTGGCTGGAGAGGAAGTACGGATATTAC
non-organic_lettuc...  GACAACTAGACCGTGGCTGGAGAGGAAGTACGGATATTAC

                       2121                                   2160
cry1A                  CATCCAAGGAGGCGATGACGTATTCAAAGAGAATTACGTT
organic_kale.........  CATCCAAGGAGGCGG-------------------------
organic_spinach......  CATCCAAGGAGGC---------------------------
organic_lettuce......  CATCCAAGGAGGCGA-------------------------
non-organic_kale.....  CATCCAAGGAGGCG--------------------------
non-organic_spinac...  CATCCAAGGAGGCGA-------------------------
non-organic_lettuc...  CATCCAAGGAGGCGA-------------------------
```

**Fig. S13. Detection of *Bacillus thuringiensis* from produce bought at a market. (A)** Standard curve of qPCR with *Bt* gDNA. The estimated equivalent number of spores were calculated by dividing the amount of gDNA in the qPCR reactions by 6.5 fg, the approximate amount of gDNA in a single *Bt* spore. **(B)** PCR gel images of plant samples. Red box indicates the *Bt* Cry1A band. The summary of *Bt* detection from produce bought from stores is shown in **Table S1**. * are negative control samples. Water was used as negative control (-) for PCR. **(C)** Alignment of PCR-amplified products from 6 produce samples showed that the PCR products are genuine *Bt* cry1A sequences. BLAST results found exclusively *Bt* in the top 100 hits. Note that the variable region (in red box) confirms the existence of 3 *Bt* variants confirming that this was not cross contamination from the lab.

Figure. S14

**A**

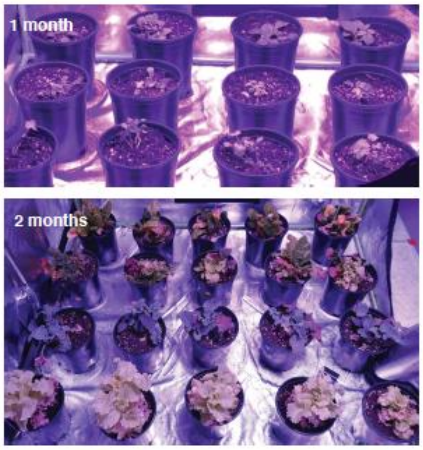


**B**

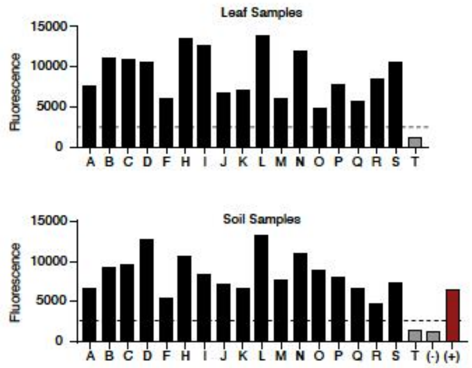


**C**

```
                Target site
ref:      ...aaGTCGACCTGCAGGCATGCAAGCtt...
plant B:  ...aaGTCGACCTGCAGGCATGCAA--tt...
plant G:  ...aaGTCGACCTGCA-GCATGCAAGCtt...
```

**D**

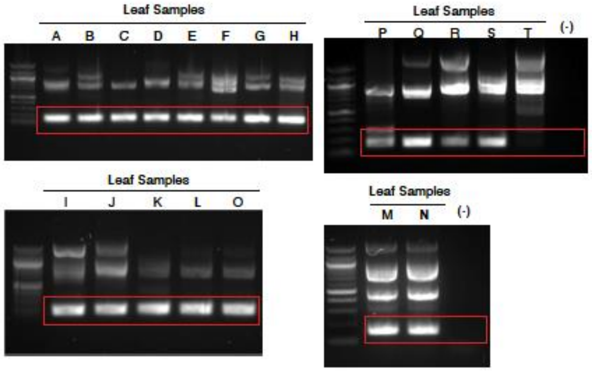


```
Plant A   TAGTACGGGCCACGATCTAAGTCGGCGG
PCR A     TAGTACGGGCCACGATCTAAGTCGG[illegible]G

Plant B   GGGATGCCCTGATAACGTAGAGTCTCAG
PCR B     GGGATGCCCTGATAACGTAGAGTCTCAG

Plant C   GAGGGCTTCTACGAAATTGCCTCACCAT
PCR C     GAGGGCTTCTACGAAATTGCCTCACCAT

Plant D   GTTCAAAAGCGGGAGTCCCGGTGAAACC
PCR D     GTTCAAAAGCGGGAGTCCCGGTGAAACC

Plant E   CTTGGTCCAATCGTATGCTAAGAGTAGC
PCR E     CTTGGTCCAATCGTATGCTAAGAGTAGC

Plant F   ACGTAGGGGGGCGCGTAACCACTAGCTC
PCR F     ACGTAGGGGGGCGCGTAACCACTAGCTC

Plant G   CCCCGTGTGGTAACTACGCAAGCCTAAC
PCR G     CCCCGTGTGGTAACTACGCAAGCCTAAC

Plant H   TGAATAAGCGCGGTCCCTAATGTTGGTG
PCR H     TGAATAAGCGCGGTCCCTAATGTTGGTG

Plant I   TCGTCAGTCGTAACTTGGGAACGCACAT
PCR I     TCGTCAGTCGTAACTTGGGAACGCACAT

Plant J   GTATGTGGGTACGATTGTAGGCTAGAAA
PCR J     GTATGTGGGTACGATTGTAGGCTAGAAA

Plant K   TCAGGGGAAACGAGTTAAGCAGAGGCAG
PCR K     TCAGGGGAAACGAGTTAAGCAGAGGCAG

Plant L   GTGAGTCCGGCCTATCACGTTTGGTAGG
PCR L     GTGAGTCCGGCCTATCACGTTTGGTAGG

Plant M   GAGTTACGGGTCAGGATCATTGCGCAGG
PCR M     GAGTTACGGGTCAGGATCATTGCGCAGG

Plant N   AGTGTCCCTTATTCTACTTTGAATTATC
PCR N     AGTGTCCCTTATTCTACTTTGAATTATC

Plant O   AGAACTACCTACGTGGCCACGCAAGCCC
PCR O     AGAACTACCTACGTGGCCACGCAAGCCC

Plant Q   ATGTTGGGACGCCAGACTGACACGCAAA
PCR Q     ATGTTGGGACGCCAGACTGACACGCAAA

Plant R   CCTACACCTTCACCGAGTGTGGAGCAAG
PCR R     CCTACACCTTCACCGAGTGTGGAGCAAG

Plant S   CTTTGGGGTTGGATAAATCTGTCGTGGT
PCR S     CTTTGGGGTTGGATAAATCTGTCGTGGT
```

Figure. S14

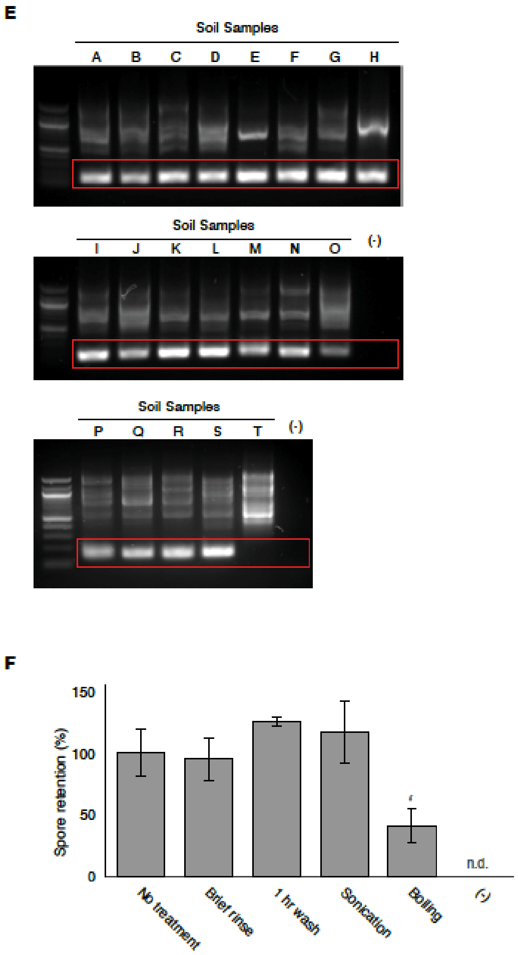


**Fig. S14. Detection of *B. subtilis* from plants grown in laboratory farm. (A)** Photographs of produce at the time of first spraying (1 month post planting) and before harvest (2 months). **(B)** SHERLOCK signals from reaction performed on leaf and soil samples using the group 2 crRNA. Plant T was not sprayed. **(C)** Plant sample E & G with alternate group barcode sequences. **(D)** PCR gel images of FMS from plant samples grown in the lab farm. Red box indicates the band that corresponds to *Bt* Cry1A. Alignment of PCR-amplified products from plant samples to sprayed FMS. **(E)** PCR gel images of FMS from the soil samples of the pots in the lab farm. **(F)** FMS retention measured by qPCR, normalized to no-treatment. FMS sprayed on plant samples were retained after rinsing, washing, sonication or boiling (see Methods). Water was used as negative control (-). n.d. = not detected.

Figure. S15

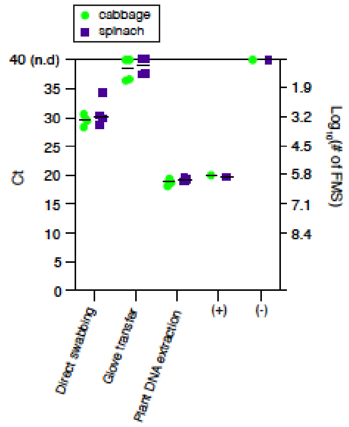


**Fig. S15. FMS remain on plant leaves.** qPCR measurement of FMS from swabbing directly the plant, swabbing from the gloves after transfer, or DNA extractions of different FMS sprayed cabbage and spinach. Non-FMS sprayed plant was used as negative control (-). n.d. = not detected.

| Organic | Sample | Merchant/Source | BT |
|---|---|---|---|
| NA | Green beans | C-Mart | - |
| NA | Okra | C-Mart | - |
| NA | Scallion | C-Mart | - |
| - | Baby spinach | Stop & Shop | + |
| - | Broccolini | Stop & Shop | - |
| - | Green beans | Stop & Shop | - |
| - | Iceberg lettuce | Stop & Shop | + |
| - | Kale | Stop & Shop | + |
| - | Okra | Stop & Shop | - |
| - | Romaine lettuce | Stop & Shop | + |
| - | Scallion | Stop & Shop | - |
| + | Baby spinach | Stop & Shop | + |
| + | Kale | Stop & Shop | + |
| + | Romaine lettuce | Stop & Shop | + |
| - | Basil | Whole Food | - |
| - | Green beans | Whole Food | - |
| - | Okra | Whole Food | - |
| + | Arugula | Whole Food | + |
| + | Baby spinach | Whole Food | + |
| + | Broccoli | Whole Food | + |
| - | Watercress | Stop & Shop | + |
| - | Baby spinach, brand B | Stop & Shop | + |
| + | Romaine lettuce, brand B | Stop & Shop | + |
| + | Cucumber | Stop & Shop | - |
| + | Green beans, brand B | Stop & Shop | - |
| - | Broccoli brand B | Stop & Shop | + |
| - | Mini peppers | Stop & Shop | + |
| - | Spinach, brand B | Stop & Shop | + |
| - | Baby arugula | Stop & Shop | + |
| - | Strawberries | Stop & Shop | - |
| - | Brussels sprouts | Stop & Shop | + |
| + | Iceberg lettuce, brand B | Stop & Shop | + |

**Table S1. Detection of *Bacillus thuringiensis* on produce samples**

Results of PCR test targeting the Cry1A gene, performed on a variety of produce samples.
+, positive -, negative, NA, not available

*For the following large tables, please see "Supplementary Tables S1-S7.xlsx"*

**Table S2. List of strains used in this study**

Bacterial and yeast strains used in this study. List includes wild type strains and mutants generated as part of this study. All unmarked *B. subtilis* mutations are in-frame deletions generated by Cre-mediated recombination and contain a lox72 scar.

**Table S3. List of primers used in this study**

List of primer sequences used in this study.

**Table S4. List of crRNA used in this study**

List of crRNA sequences used for SHERLOCK reactions in this study.

**Table S5. List of barcodes used in this study**

List of unique barcode sequences used in this study, see Fig. S1 for more detailed barcode design.

**Table S6. List of synthetic megamers used in this study**

List of synthetic megamer sequences used for SHERLOCK reactions in this study.

| Surface Material | Count | Control Conditions | Perturbation Conditions | Direct Sample Technique | Transfer Sample Technique |
|---|---|---|---|---|---|
| Sand | 3 control & 3 perturbed | **Temperature:** 25 C Fixed<br>**Wind:** None<br>**Rain:** None | **Temperature:** Weekly oscillation (25-35C)<br>**Wind:** Fans, fixed speed<br>**Rain:** weekly, varying intensity from spraying to watering can | Powersoil | Object Swab, NaOH lysis |
| Soil | 3 control & 3 perturbed | **Temperature:** 25 C Fixed<br>**Wind:** None<br>**Rain:** None | **Temperature:** Weekly oscillation (25-35C)<br>**Wind:** Fans, fixed speed<br>**Rain:** weekly, varying intensity from spraying to watering can | Powersoil | Object Swab, NaOH lysis |
| Carpet | 2 control & 2 perturbed | **Temperature:** 25 C Fixed<br>**Humidity:** 40-50% Controlled<br>**Cleaning:** None | **Temperature:** 25 C Fixed<br>**Humidity:** 40-50% Controlled<br>**Cleaning:** Vacuuming | Surface Swab, NaOH lysis | Object Swab, NaOH lysis |
| Wood | 2 control & 2 perturbed | **Temperature:** 25 C Fixed<br>**Humidity:** 40-50% Controlled<br>**Cleaning:** None | **Temperature:** 25 C Fixed<br>**Humidity:** 40-50% Controlled<br>**Cleaning:** Sweeping | Surface Swab, NaOH lysis | Object Swab, NaOH lysis |

**Table S7. Description of incubator experiment**

Description matrix of conditions used in the incubator experiment, including control conditions, perturbations, and sampling techniques.
